# Mechanistic basis for nigericin-induced NLRP1 inflammasome activation in human epithelial cells

**DOI:** 10.1101/2023.06.23.546021

**Authors:** Pritisha Rozario, Miriam Pinilla, Anna Constance Vind, Kim S. Robinson, Toh Gee Ann, Muhammad Jasrie Firdaus, José Francisco Martínez, Lin Zhewang, Simon Bekker-Jensen, Etienne Meunier, Franklin Zhong

## Abstract

Nigericin, an ionophore derived from *Streptomyces hygroscopicus*, is arguably the most commonly used tool compound to study the NLRP3 inflammasome. Recent findings, however, showed that nigericin also activates the NLRP1 inflammasome in human keratinocytes. In this study, we resolve the mechanistic basis of nigericin-driven NLRP1 inflammasome activation. In multiple non-hematopoietic cell types, nigericin rapidly and specifically inhibits the elongation stage of the ribosome cycle by depleting cytosolic potassium ions. This activates the ribotoxic stress response (RSR) sensor kinase ZAKɑ, p38 and JNK, as well as the hyperphosphorylation of the NLRP1 linker domain. As a result, nigericin-induced pyroptosis in human keratinocytes is blocked by extracellular potassium supplementation, ZAKɑ knockout or pharmacologic inhibitors of ZAKɑ and p38 kinase activities. By surveying a diverse panel of ionophores, we show that the electroneutrality of potassium efflux is essential to activate ZAKɑ-driven RSR, likely because a greater extent of K+ depletion is necessary to activate ZAKɑ-NLRP1 than NLRP3. These findings resolve the mechanism by which nigericin activates NLRP1 in nonhematopoietic cell types and demonstrate an unexpected connection between RSR, perturbations of potassium ion flux and innate immunity.

**GRAPHICAL ABSTRACT:** 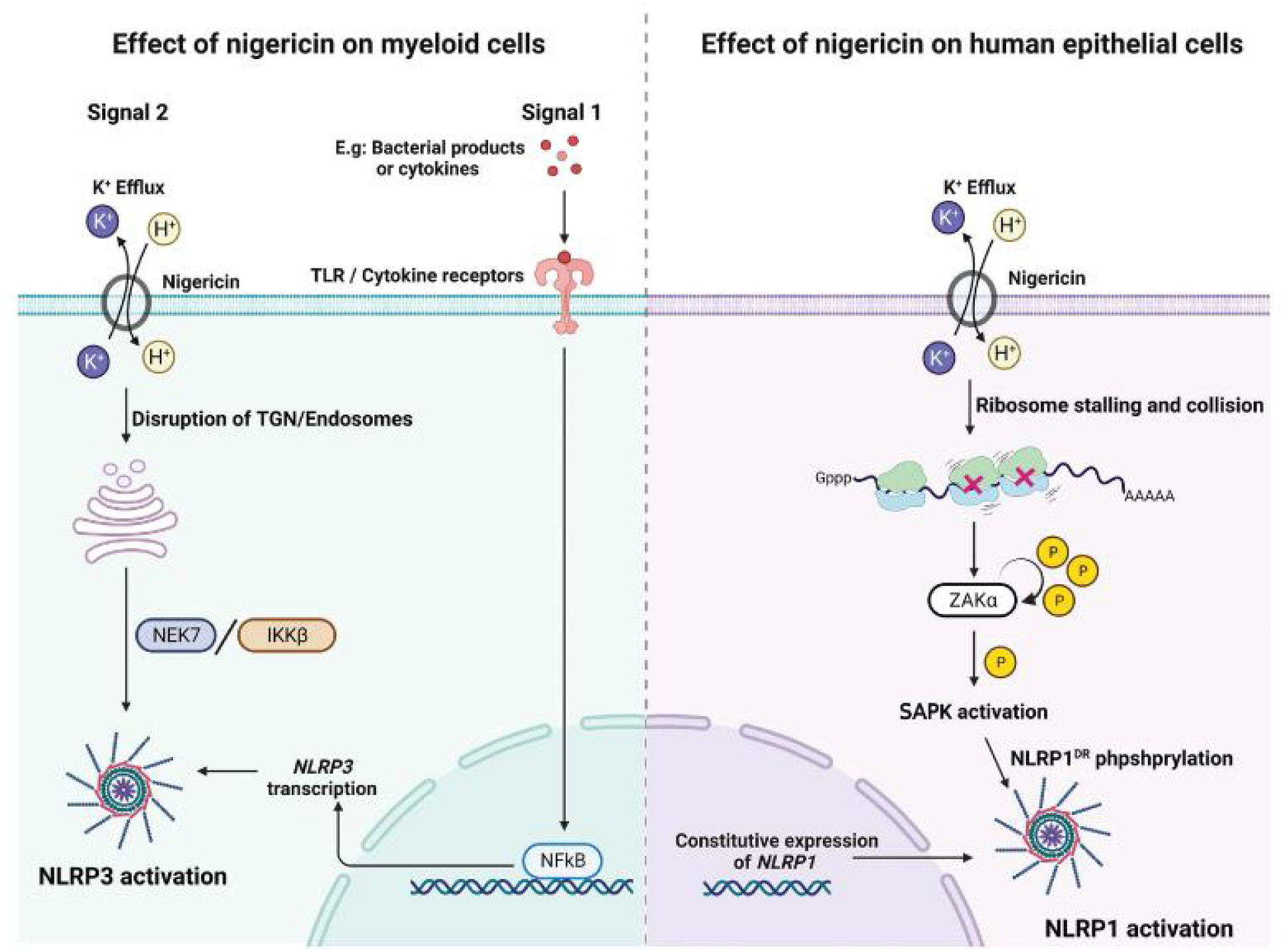

**SIGNIFICANCE:** Nigericin is familiar to the inflammasome field as the most robust and commonly used NLRP3 inducer. It has enabled numerous breakthroughs in the field linking NLRP3 activation to potassium efflux. In this manuscript, we report that nigericin activates an alternate inflammasome sensor, NLRP1 in primary human skin, nasal and corneal epithelial cells. NLRP1 activation by nigericin requires K+ efflux-driven ribosome stalling and the ribotoxic stress response (RSR) sensor MAP3K, ZAKɑ. We further identify the key biophysical principles that explain why only a subset of K+ ionophores, exemplified by nigericin, function as ‘super’ inflammasome agonists that can activate either NLRP1 or NLRP3, depending on cell type. These results reveal an unexpected connection between RSR, potassium ion flux and innate immunity.

## INTRODUCTION

It has been estimated that mammalian cells consume up to ∼70% of the total energy expenditure actively transporting sodium and potassium ions across the plasma membrane, creating a high potassium (∼140mM) and low sodium (∼5-15mM) environment in the cytosol relative to the extracellular space ^1^. The resulting electrochemical potential across the plasma membrane is essential to many physiological processes, e.g. intracellular uptake of nutrients, transmission of neuronal action potential and contraction of muscles among others. Dysregulation of membrane potential and ion flux, cause a variety of human diseases, including neurological disorders, cystic fibrosis and heart failure ^2, 3^.

Ionophores are molecules that facilitate the transport of ions across lipid bilayers. As a result, ionophores are widely used as chemical probes to study the biological function of ionic flux, or as pharmacologic agents to manipulate ionic homeostasis in vivo ^4, 5^. Nigericin (CAS 28643-80-3), a secondary metabolite isolated from *Streptomyces hygroscopicus*, is a well studied carrier-type ionophore that specifically ‘anti-ports’ potassium ions in exchange for protons. This process allows both ions to flow down their electrochemical gradients without incurring a net electrical charge (electroneutral) and is therefore thermodynamically favorable. As a result, nigericin is capable of completely dissipating the K+ and proton gradients where they run in opposite directions, e.g. across the plasma membrane of mammalian cells. Due to this property, nigericin has been widely used in chemistry as a K+ or proton ‘clamp’ to ensure stable equalization of the respective ionic concentrations across diffusion barrier membranes, or to probe the consequences of K+ movements in biological systems ^6^.

The plasma membrane of eukaryotic cells is prone to damage caused by environmental toxins, endogenous danger signals and microbial infections, all of which can lead to the leakage of cytosolic K+ ions into the extracellular space. In vertebrate species, K+ efflux is specifically sensed by a conserved intracellular innate sensor protein known as NLRP3 ^7–12^. The depletion of cytosolic potassium ions below a certain concentration, which could be caused by ionophores, pore-forming toxins and ATP, causes NLRP3 to rapidly oligomerize and assemble the ‘inflammasome’ complex, which in turns activates pro-caspase-1 via the adaptor protein ASC. This process induces rapid pyroptotic cell death via the pore-forming protein GSDMD and the secretion of pro-inflammatory cytokines in the IL-1 family. Due to its ability to rapidly trigger K+ efflux, nigericin is arguably the most widely used agonist to probe NLRP3 function. For instance, nigericin was instrumental in enabling the elucidation of the atomic structure of activated NLRP3 ^11, 13–15^. It was also used to uncover the critical involvement of trans-Golgi and vesicular lipid molecules in NLRP3 function ^16–18^. It is generally thought that nigericin is an exclusive NLRP3 agonist, at least in hematopoietic cell types, as numerous studies demonstrated that chemical inhibition or genetic ablation of NLRP3 completely abrogates nigericin-driven pyroptosis in human and murine macrophages and monocytes, while other inflammasome sensor proteins, such as NLRC4, AIM2 and MEFV^9, 19^, which are also expressed in cells of the myeloid origin, do not respond to nigericin. As such, nigericin-driven pyroptosis is often used as a diagnostic test for NLRP3 functionality in various experimental systems.

Notwithstanding the indisputable evidence showing that nigericin activates NLRP3, some studies suggest that nigericin has NLRP3-independent functions in programmed cell death. For instance, nigericin is cytotoxic in many human cancer cell lines with IC50 ∼5µM, including those that do not express inflammasome components ^20^. Recent studies additionally demonstrated that nigericin can induce pyroptosis in human keratinocytes via another inflammasome sensor protein known as NLRP1 ^21, 22^, which is predominantly expressed in human skin and airway and does not share the same agonist repertoire as NLRP3 ^23, 24^. These findings prompted us to re-examine the role of nigericin outside the context of NLRP3 agonism, and search for novel links between K+ efflux and human innate immunity. Here, we report that nigericin-induced K+ efflux inhibits protein synthesis due to a specific block of the ribosomal elongation cycle. By doing so, nigericin functions as a potent inducer of the ZAKɑ-dependent ribotoxic stress response (RSR) in numerous human cell lines, and NLRP1 phosphorylation more specifically in primary human epithelial cells. By surveying a range of ionophores, we report the electroneutrality of K+ efflux is essential for RSR induction. This explains why only K+/H+ antiporters, exemplified by nigericin and the veterinary antibiotic lasalocid acid ^8, 25^, are capable of activating both NLRP1 and NLRP3. Our results shed light on how nigericin activates the NLRP1 inflammasome in non-hematopoietic cell types, and suggests that RSR and the NLRP1 inflammasome have a conserved function in the innate immune defense against perturbations of ionic potential.

## RESULTS

### Nigericin-driven pyroptosis in human epithelial cells depends on NLRP1 and potassium efflux

We first sought to validate the findings published by Fenini et al demonstrating that nigericin activates NLRP1 ^21^. In independently generated primary keratinocytes derived from healthy donor biopsies, nigericin indeed caused cardinal features of pyroptosis in a dose dependent manner, as evidenced by membrane swelling (Figure 1A), GSDMD cleavage (Figure 1B), ASC speck formation (Figure 1C, S1F), rapid propidium iodide (PI) uptake (Figure S1A-B) and IL-1 secretion (Figure 1B, Figure S1C). This effect was rapid (detectable within 3 hours) and required similar or lower concentrations (1-5 µM) typically used to activate NLRP3 (Figure S1B-C). Importantly, nigericin-driven inflammasome assembly and pyroptosis are abrogated by caspase-1 inhibitor belnacasan (Figure S1E) or CRISPR/Cas9 KO of *NLRP1* in primary keratinocytes (Figure 1D-E, S2C), but is insensitive to the NLRP3 inhibitor MCC950 (Figure S2A-B). This is in contrast to NLRP3-expressing THP1 or U937 cells where nigericin-triggered IL-1β is completely abrogated by MCC950 (Figure S3A-B). Non-ionophore NLRP3 agonist ATP fails to activate pyroptosis in keratinocytes up to 1 mg/mL (Figure S1D). In addition to skin keratinocytes, nigericin was also able to activate IL-18 secretion in primary human nasal epithelial cells (HNECs) and corneal epithelial cells (HCEs) in a NLRP1-dependent manner (Figure 1F-G), and ASC-GFP speck formation in a A549-ASC-GFP reporter cell line where NLRP1 is exogenously expressed (Figure S2D). These results confirm that nigericin indeed activates both endogenously expressed and reconstituted NLRP1 inflammasome in epithelial cells. Curiously, we found that the immortalized keratinocyte cell line N/TERT-1 (also referred as N/TERT), which has increasingly been used to study the NLRP1 inflammasome, has diminished ability to undergo NLRP1-driven pyroptosis when treated with nigericin (Figure S3C). These results are consistent with recent findings that N/TERT cells also have reduced sensitivity to bacterial ribotoxins such as DT and exoA ^26^. While the exact reasons are unclear, this difference underscores the importance of using primary cells in investigating NLRP1 function *in vitro*. In all subsequent experiments, the effect of nigericin on pyroptosis was studied exclusively in primary cells, while N/TERT cells were used to study the upstream molecular mechanisms.

**Figure 1.**
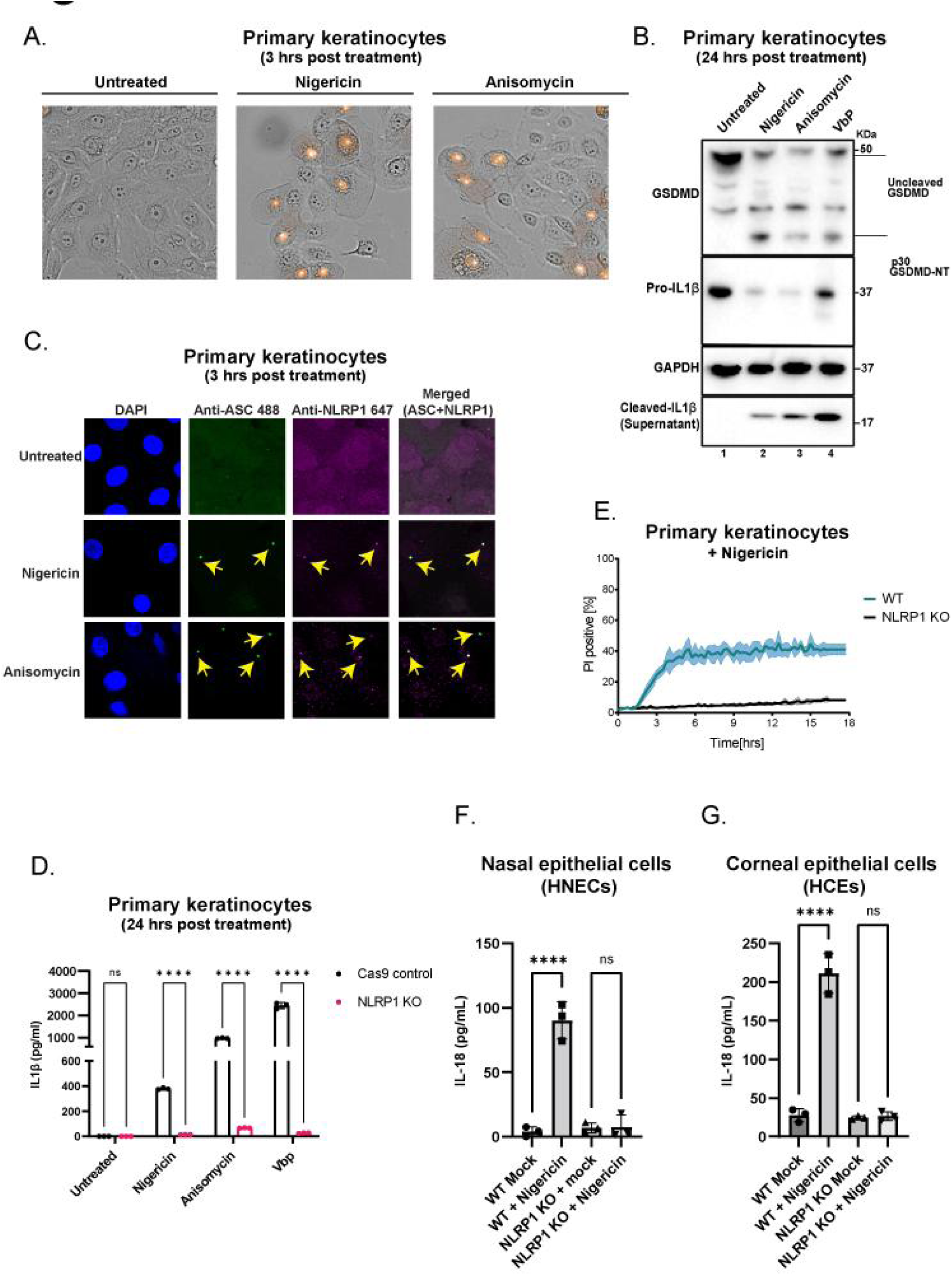
A. Representative bright-field microscopy images of primary keratinocytes showing propidium iodide (PI) inclusion 3 hours post treatment with nigericin (5µg/mL) or anisomycin (1µM). B. Western blot analysis of primary keratinocytes showing GSDMD (full length and cleaved), pro-IL1β, cleaved IL1β and GAPDH (loading control) upon overnight treatment with nigericin (5µg/mL), anisomycin (1µM) or VbP (3µM). Immunoblot representative of three replicate experiments. C. Representative confocal microscopy images showing ASC (green), NLRP1 (far-red) and DAPI (blue) staining in primary keratinocytes 3 hours post treatment with nigericin (5µg/mL) or anisomycin (1µM). Yellow arrows indicate speck formation by ASC and NLRP1. D. ELISA showing IL-1β secretion in primary keratinocytes after nigericin (5µg/mL), anisomycin (1µM) or VbP (3µM) treatment. Supernatant was harvested 24 hours post treatment. E. Quantification of the percentage of PI positive WT and NLRP1 KO primary keratinocytes after nigericin (5µg/mL) treatment. Cells were imaged at 15 minute intervals for 18 hours. F. ELISA showing IL-18 secretion in WT and NLRP1 KO human nasal epithelial cells (HNECs) after nigericin (5µM) treatment. Supernatant was harvested 24 hours post treatment. G. ELISA showing IL-18 secretion in WT and NLRP1 KO human corneal epithelial cells (HCEs) after nigericin (5µM) treatment. Supernatant was harvested 24 hours post treatment.

Next we tested if nigericin-dependent NLRP1 activation depends on K+ efflux or represents an ionophore independent role of nigericin. In primary keratinocytes, supplementing the culture media with 50 mM KCl abrogated the rapid PI uptake (Figure 2A, S4A) and other hallmarks of pyroptosis induced by nigericin, including GSDMD cleavage and IL-1β secretion (Figure 2C-D,S4B). By contrast, 50mM KCl did not alter the kinetics and extent of PI uptake following anisomycin (ANS) treatment (Figure 2B). Unlike extracellular KCl, the cell permeable Ca^2+^ chelator BAPTA-AM did not significantly reduce IL-1β secretion caused by nigericin (Figure 2E). Although 50mM KCl also had a mild inhibitory effect on VbP- and ANS-driven IL-1β secretion, this could be mostly accounted for by reduced levels of uncleaved pro-IL-1β (Figure 2D). These data support the notion that similar to NLRP3, nigericin activates the NLRP1 inflammasome specifically via K+ efflux.

**Figure 2.**
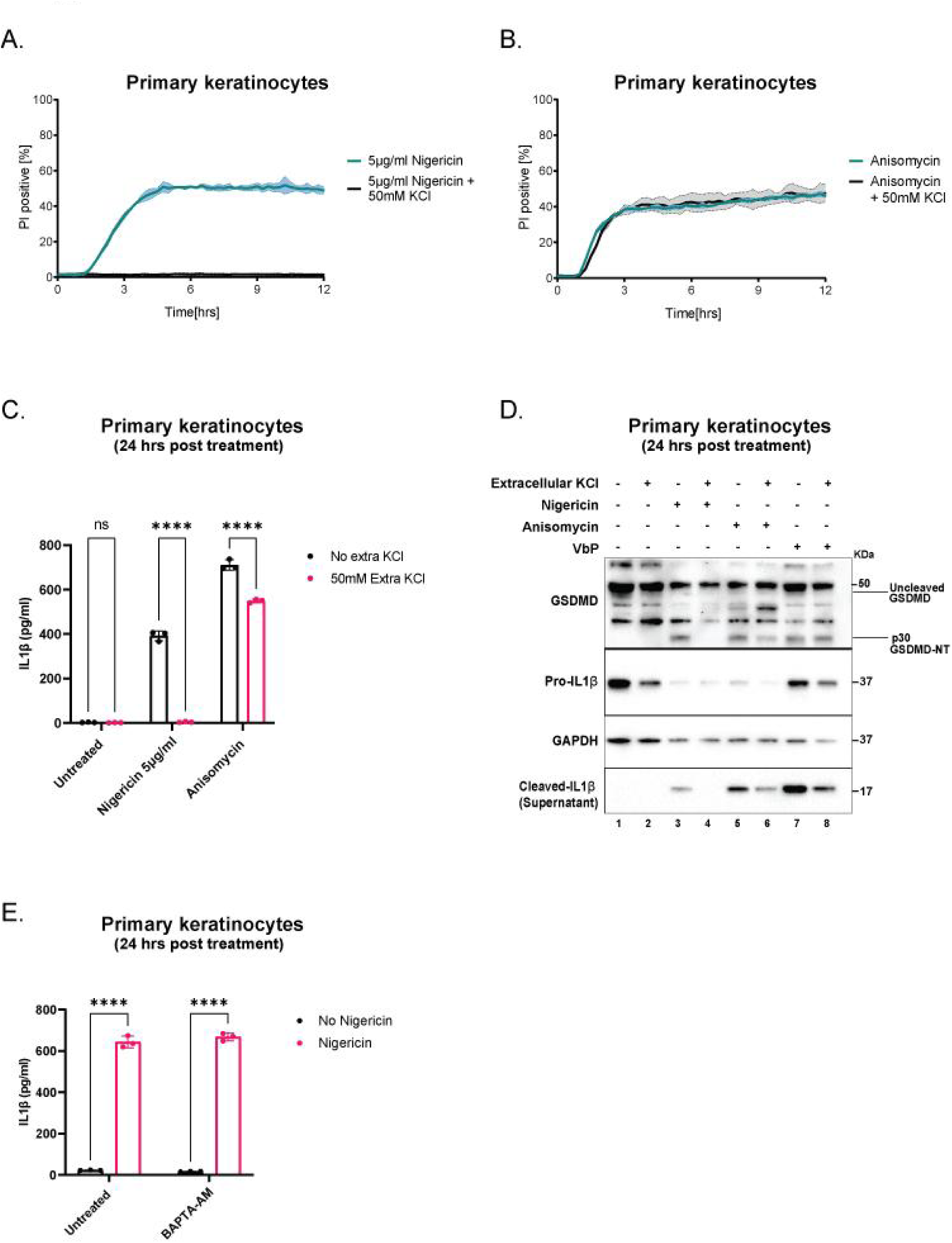
A. Quantification of the percentage of PI positive primary keratinocytes treated with nigericin (5µg/mL) with or without 50mM KCl. B. Quantification of the percentage of PI positive primary keratinocytes treated with anisomycin (1µM) with or without extracellular KCl. Cells were imaged at 15 minute intervals for 12 hours. C. ELISA showing IL-1β secretion from media in (A) and (B). D. Western blot analysis of WT primary keratinocytes showing GSDMD (full length and cleaved), pro-IL1β, cleaved IL1β and GAPDH after overnight treatment with nigericin (5µg/mL), anisomycin (1µM) or VbP (3µM) with or without extracellular KCl (50mM). Immunoblot representative of three replicate experiments. E. ELISA showing IL-1β secretion from supernatant obtained from WT primary keratinocytes following a 15 min pre-treatment with BAPTA-AM (5µM) and overnight treatment with nigericin (5µg/mL).

### Nigericin blocks protein synthesis and induces ribotoxic stress (RSR) via potassium efflux

Even though NLRP3 and NLRP1 are generally not expressed in the same cell types ^11, 17^,it is conceptually conceivable that potassium efflux activates both sensors via the same mechanism. However, unlike NLRP3, activated NLRP1 localizes almost exclusively to ASC specks and not to any obvious organellar or endosomal structures (Figure 1C). Recently, we and others reported that NLRP1 can be activated by the ribotoxic stress response kinase ZAKɑ downstream of ribosome stalling and/or collisions ^26–29^. This discovery explains why NLRP1 can sense seemingly unrelated triggers, including UVB irradiation, small molecule ribosome inhibitors such as ANS and at least three known bacterial exotoxins. To the best of our knowledge, nigericin has not been linked to RSR, but several lines of evidence support this connection. First, nigericin and other NLRP3 agonists have been shown to inhibit protein synthesis in murine bone marrow derived macrophages ^30^. In addition, in erythroid cell lysates in vitro, the rates of peptidyl transfer reaction by the ribosome are sensitive to potassium ion concentration ^31^. More importantly, updated atomic structures of the human ribosome complex revealed an abundance of potassium ions near the peptidyl transfer center ^32^. Based on these results, we hypothesized that nigericin activates NLRP1 via ZAKɑ-driven RSR. Indeed, 3 hours of nigericin treatment led to a near complete cessation of de novo protein synthesis in primary keratinocytes, as revealed by the extent of puromycin incorporated into the nascent polypeptides (Figure 3A-B). In addition, the effect of nigericin on translation requires the integrity of the plasma membrane, as the translation of a synthetic mRNA in rabbit reticulocyte lysates (RRL) is not inhibited, and even slightly enhanced, by nigericin (Figure 3A, C). Furthermore, extracellular K+ at 50mM restored protein synthesis in nigericin-treated cells (Figure 3D, lane 1 vs lane 8). Taken together, these data suggest that nigericin inhibits translation indirectly by depleting K+ from the cytosol, rather than acting as a direct competitive/allosteric inhibitor of the ribosome.

**Figure 3.**
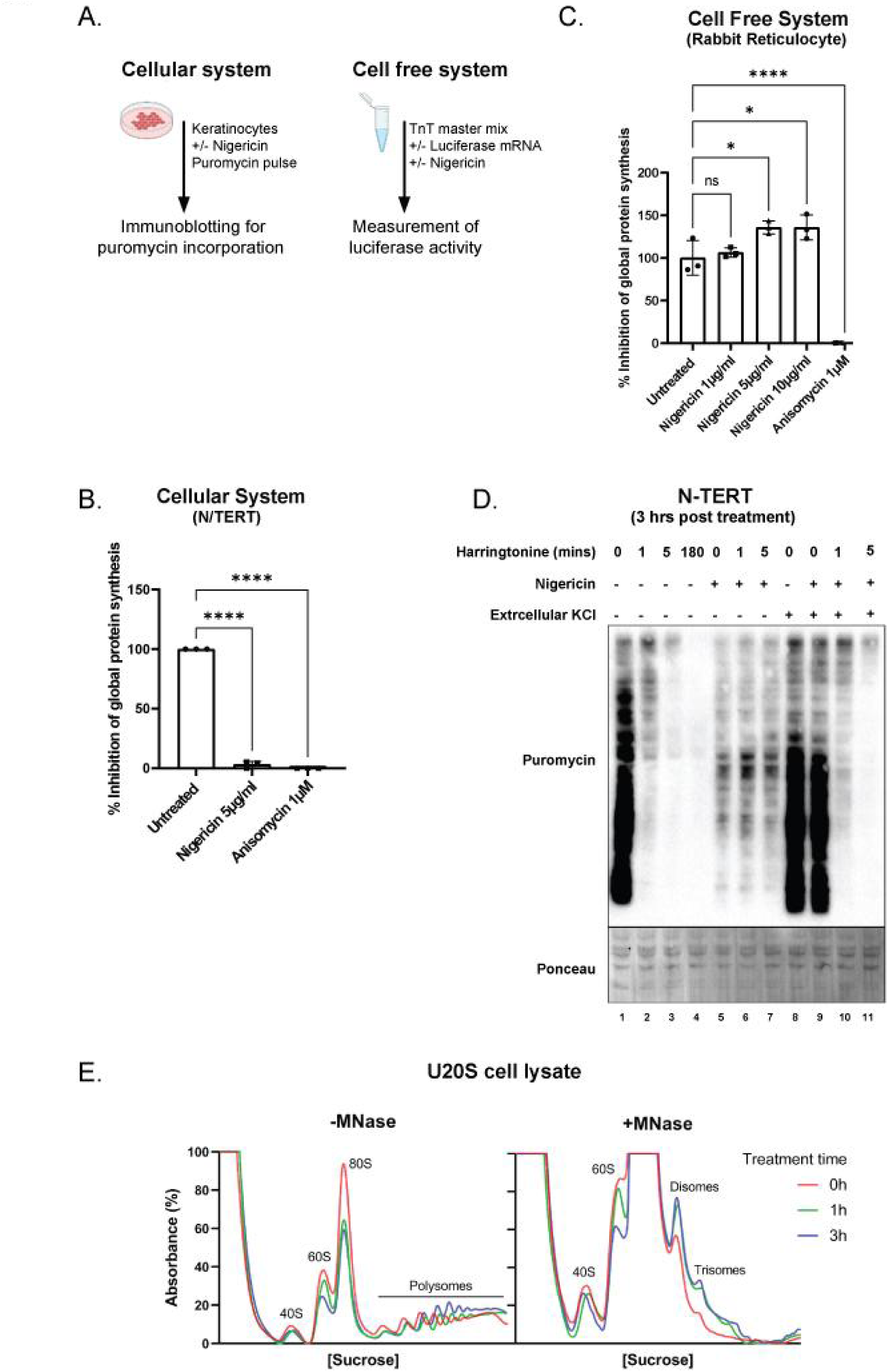
A. Experimental setup to measure the effect of nigericin in cellular and cell free systems. B. Quantification of the percentage inhibition of global protein synthesis in the cellular system (N/TERT) following 3 hour treatment with nigericin (5µg/mL) or anisomycin (1µM). Protein synthesis was measured by normalized lane intensity in anti-puromycin blots from three independent biological replicates. Percentage was calculated relative to untreated cells. C. Quantification of the percentage inhibition of in vitro protein synthesis in the cell free system (rabbit reticulocyte) following 1 hour treatment with nigericin (1µg/mL, 5µg/mL or 10µg/mL) or anisomycin (1µM). Protein synthesis was measured by luciferase activity. Percentage was calculated relative to untreated lysate. D. Western blot analysis of N/TERT cells subjected to the harringtonine run-off assay. Cells were treated with nigericin (5µg/mL) with or without KCl (50mM) for 3 hours before the addition of harringtonine (2µg/mL) for the respective durations. Runoff was terminated by puromycin (10µg/mL) for 10 mins. Ponceau staining was used as the loading control. Immunoblot representative of three replicate experiments. E. Polysome profiles from undigested (left) or MNase-digested (right) U2OS cell lysates after 1 hour nigericin (10 µg/mL) (green) or 3 hour nigericin treatment (blue).

To investigate which stage of the ribosome cycle is inhibited by nigericin-induced K+ efflux, we carried out a harringtonine run-off assay in N/TERT cells (Figure S5A), where the amount of puromycin-labeled nascent peptides is measured at regular intervals after the translational inhibition is blocked by harringtonine. In this assay, the time-dependent decrease in puromycin incorporation, i.e. the ‘run-off’ rate, is directly proportional to the speed of ribosome elongation ^33^. In mock treated cells, most of the ribosomes completed the ‘run-off’ within 5 mins. By contrast, in nigericin-treated cells, the level of puromycin-labeled proteins remained constant across time, suggesting that the majority of the 80S ribosomes became arrested (Figure 3D). Supplementing culture media with 50mM KCl restored the ribosome elongation to levels that are similar to untreated controls (Figure 3D). Taken together, these results demonstrate that K+ efflux blocks the elongation stage of the ribosome cycle, resulting in ribosome stalling. It is critical to note that all aforementioned puromycin-incorporation assays (Figure 2B, 2D) were carried out in the presence of the pan-caspase inhibitor, emricasan, which blocks cell death including caspase-1 driven pyroptosis. Thus, nigericin-driven ribotoxic stress occurs upstream, rather than downstream of cell death.

We next tested the effect of nigericin on U2OS cells, which do not express any inflammasome components but retain a fully functional RSR pathway ^34, 35^. Nigericin also causes ribosome stalling in a dose dependent manner in U2OS cells, as evident in the increased amounts of polysomes at the expense of monosomes in the polysome gradient (Figure 3E - left). To understand the mechanistic basis of nigericin-induced ZAKa activation, we looked for the presence of ribosome collisions. By digesting lysates from nigericin-treated cells with MNase before resolving the polysomes on a sucrose gradient, we could detect the formation of “disomes” (Figure 3E - right), which are MNase resistant polysomes peaks indicating the presence of collided ribosomes. This was supported by the appearance of collision markers such as EDF1 recruitment and RPS10 ubiquitination after nigericin treatment (Figure S5B). Thus, nigericin causes translational arrest by stalling elongating ribosomes in multiple cell types. Although we cannot rule out the possibility that nigericin might affect other aspects of translation such as ribosome assembly, these effects were not apparent in the polysome gradient analysis, as the 40S, 60S and 80S peaks were still clearly distinguishable (Figure 3E - left). Thus, nigericin preferentially inhibits the elongation stage of the ribosome cycle, as predicted by previous in vitro and structural studies ^31, 32^.

### ZAKɑ is required for nigericin-driven pyroptosis in non-hematopoietic cells

We and others recently characterized the biochemical mechanisms by which a ribosome-bound kinase, ZAKɑ activates ribotoxic stress signaling in response to ribosome stalling/collisions ^34–37^. For unknown reasons, ribosome elongation inhibitors differ markedly in their abilities to activate ZAKɑ in a manner that is not correlated with their abilities to block protein synthesis. For instance, emetine and cycloheximide, are poor activators of ZAKɑ despite being strong inhibitors of global protein synthesis; whereas ANS and hygromycin act as strong activators of ZAKɑ ^34, 35^. In primary keratinocytes, we found that nigericin belongs to the latter group, as it strongly induces ZAKɑ auto-phosphorylation (Figure 4A) and MAP2K-driven p38 and JNK phosphorylation in a K+ efflux-dependent manner (Figure 4A, Figure B, lanes 4-6 vs 7-9). Using a live cell imaging based p38 and JNK Kinase Translocation Reporter (KTR) system, we found that nigericin-induced JNK and p38 activation occurs within 10 mins in N/TERT cells, similar to ANS (Figure 4C-D, Figures S6A-B). ZAKɑ KO also eliminated JNK and p38 phosphorylation in nigericin-treated U2OS and HeLa cells (Figure S7A-B). These findings corroborate that nigericin triggers bona fide ZAKɑ-driven RSR in a K+-efflux dependent manner in a variety of cell types, including those that do not express NLRP1.

**Figure 4.**
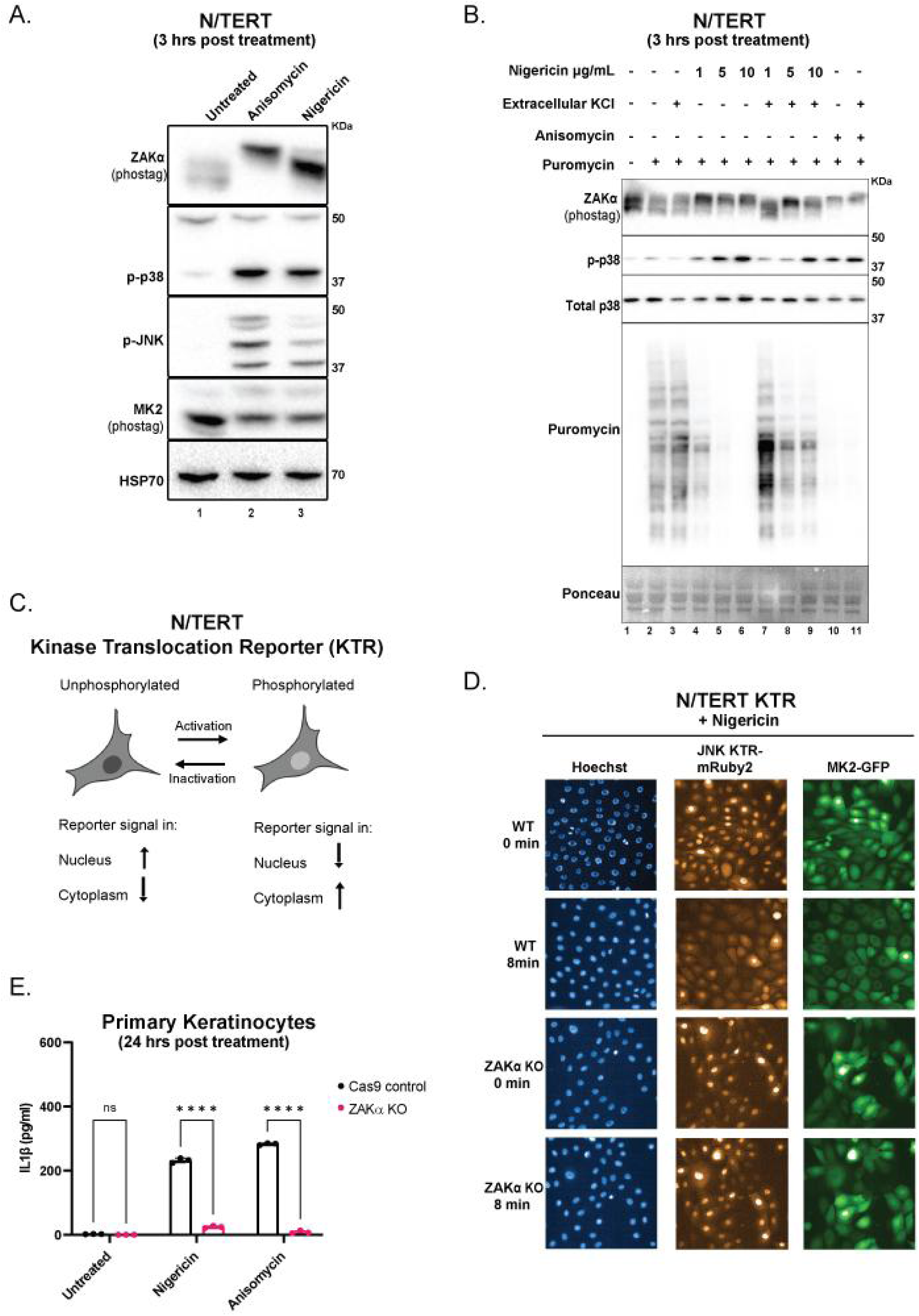
A. Western blot analysis of PhosTag SDS-page (ZAKɑ and MK2) or SDS-PAGE (p-p38 and p-JNK) in N/TERT cells after nigericin (5µg/mL) or anisomycin (1µM) treatment. Immunoblot representative of three replicate experiments. B. Western blot analysis of PhosTag SDS-PAGE (ZAKɑ) or SDS-PAGE (p-p38, total p38 and puromycin) in N/TERT cells upon 3 hour treatment with nigericin (5µg/mL) or anisomycin (1µM) with or without supplementation of extracellular KCl (50mM). Ponceau staining was used as the loading control. Immunoblot representative of three replicate experiments. C. Schematic illustrating the kinase translocation reporter (KTR) system. D. Representative images of WT and ZAKɑ KO N/TERT cells stably expressing MK2-mEGFP and JNK KTR-mRuby2 upon stimulation with nigericin (5µg/mL). Images were acquired every 4 minutes for 2 hours. E. ELISA showing IL-1β secretion from supernatant obtained from WT or ZAKɑ KO primary keratinocytes following overnight treatment with nigericin (5µg/mL) or anisomycin (1µM).

Next we tested the requirements of ribosome stalling and ZAKɑ in the activation of the NLRP1 inflammasome by nigericin in primary cells. In the first test, we took advantage of the fact that a high dose of certain ribosome elongation inhibitors have an inverse dose response with regard to RSR induction, ie. a higher dose can block ribosome elongation without activating RSR ^35^. This is exemplified by the effect of emetine in primary keratinocytes. Emetine itself can cause RSR at 10 µM, but not at 50 µM, as assayed by p38 and JNK phosphorylation (Figure S8A-B). Pretreating primary keratinocytes with 50 µM Emetine blocked nigericin- and ANS- induced ZAKɑ auto-phosphorylation and SAPK (p38/JNK) phosphorylation (Figure S8A-B). 50 µM emetine also abrogated nigericin-induced IL-1β secretion (Figure S8C). This epistatic analysis suggests that it is not ribosome stalling per se, but rather the induction of RSR that serves as the upstream signal for nigericin-driven (as well as ANS-driven) RSR in primary keratinocytes.

In the second test, we generated ZAKɑ KO primary keratinocytes and A549-ASC-GFP-NLRP1 reporter cells. In contrast to wild-type controls, nigericin failed to induce detectable IL-1β secretion and rapid PI uptake in ZAKɑ KO keratinocytes (Figure 4E, S9A) or ASC-GFP speck formation in ZAKɑ KO A549-ASC-GFP-NLRP1 cells (Figure S10A-B). In addition, nigericin-driven NLRP1 activation can be inhibited by the same inhibitors as ANS-driven NLRP1 activation, including ZAKɑ inhibitors M443, compound 6p and p38 inhibitor Neflamapimod (Figure S9B-C). In contrast, ZAKɑ inhibition did not affect nigericin-dependent NLRP3 activation in THP1 cells, which is completely inhibited by MCC950 (Figure S11A-B). Taken together, these results establish that nigericin activates NLRP1 exclusively through ZAKɑ in human epithelial cells.

### Only electroneutral, K+ selective ionophores, exemplified by nigericin and lasalocid acid activate NLRP1

Aside from nigericin, many molecules are known to activate NLRP3 by directly or indirectly inducing K+ efflux from the cytosol. We thus investigated whether ZAKɑ and NLRP1 can also be triggered by such a wide range of K+ efflux agents. To this end, we screened a panel of naturally occurring or synthetic ionophores or molecules with ionophore-like activities, including many that have previously been shown to activate NLRP3 ^8, 9, 11^ (Table 1). All compounds were screened as equivalent concentrations up to 10 µM. The behaviors fell into three broad categories. Category I, exemplified by nigericin and lasalocid acid (an veterinary antibiotic), are electroneutral ionophores that selectively ‘antiport’ K+ and H+, i.e. without causing charge separation across the plasma membrane. Thus, these molecules are known to completely equalize the K+ concentration between the cytosol and extracellular space. Both nigericin and lasalocid acid inhibit protein synthesis and cause ribosome stalling, as measured by puromycin incorporation and ribosome runoff (Figure 5B), as well as full-blown RSR induction marked by ZAKɑ phosphorylation (Figure 5A) and SAPK activation (Figure 5A). Similar to nigericin, lasalocid causes inflammasome-driven pyroptosis in primary keratinocytes (Figure 6A-C, Figure S13C-D, S14A-B), which was abrogated by NLRP1 KO (Figure 6B, Figure S13C-D) and ZAKɑ KO (Figure S14A) or inhibition (Figure 6D). ZAKɑ KO also significantly, albeit partially, inhibited p38/JNK phosphorylation and p38/JNK KTR translocation triggered by lasalocid acid (Figure S12B, S13A), suggesting that it can trigger both ZAKɑ-driven RSR and ZAKɑ-independent stress signaling. In support for this notion, lasalocid was found to induce GSDME cleavage in an ZAKɑ-independent manner (Figure S14A, lane 11-12).

**Figure 5.**
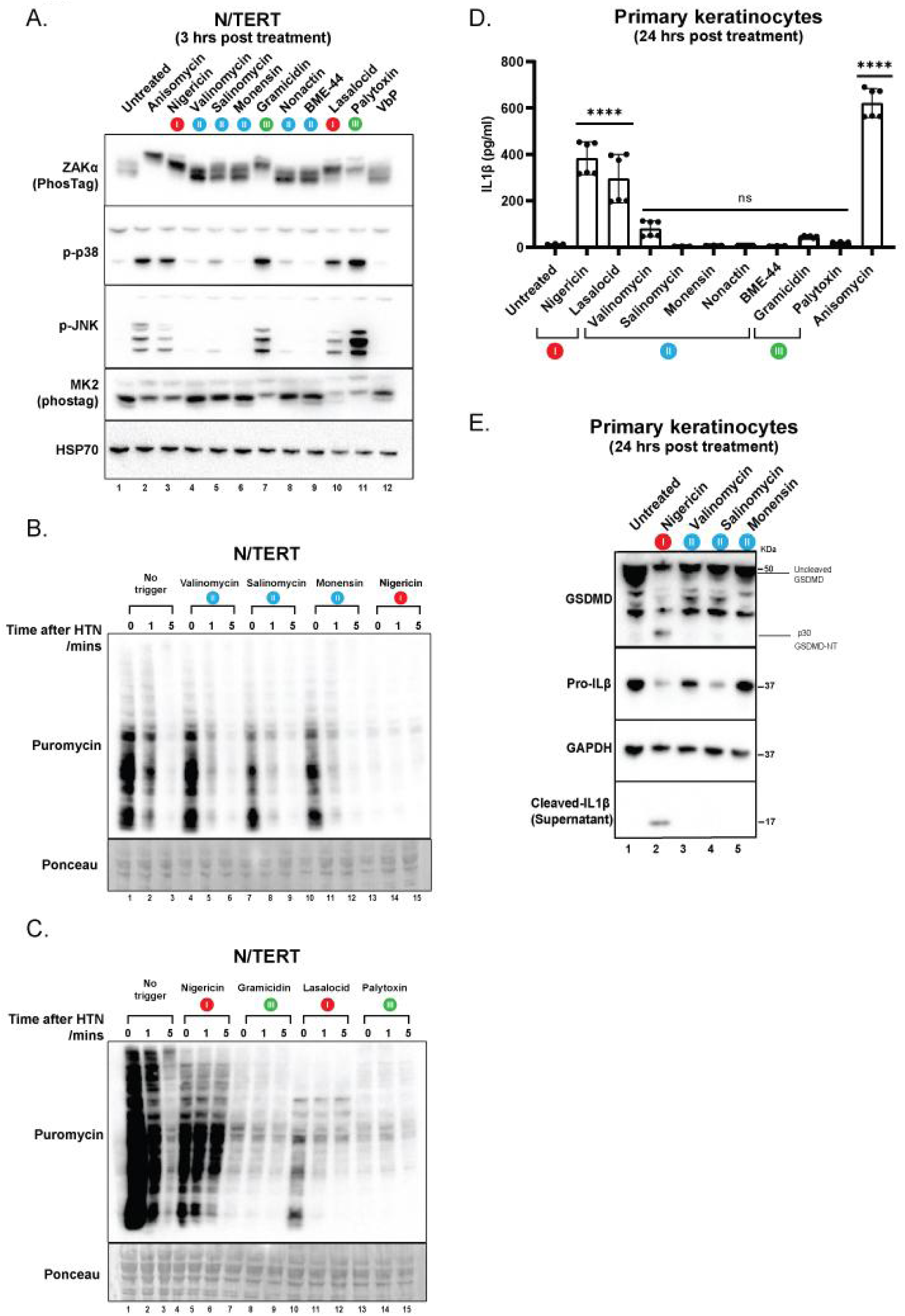
A. Western blot of PhosTag SDS-PAGE (ZAKɑ and MK2) or SDS-PAGE (p-p38,p-JNK and HSP70) N/TERT cells treated with anisomycin (1µM), nigericin (5µg/mL), valinomycin (5µM), salinomycin (5µM), monensin (5µM), gramicidin (1µM), nonactin (5µM), BME-44 (5µM), lasalocid (5µM), palytoxin (100pM) or VbP (3µM). Data are representative of three independent experiments. Immunoblot representative of three replicate experiments. B. Anti-puromycin immunoblots of N/TERT cells subjected to harringtonine run-off. Cells were treated for 3 hours with valinomycin (5µM), salinomycin (5µM), monensin (5µM), nigericin (5µg/mL) before harringtonine addition (2µg/mL) for the respective durations. Nascent peptides were then labeled with puromycin (10µg/mL) for 10 mins. Ponceau staining was used as the loading control. Immunoblot representative of three replicate experiments. C. Same as (B) except cells were treated with gramicidin (1µM), lasalocid (5µM) or palytoxin (100pM). D. ELISA showing IL-1β secretion in primary keratinocytes following overnight treatment with various potassium ionophores and anisomycin. E. Western blot analysis of primary keratinocytes showing GSDMD (full length and cleaved), pro-IL1β, cleaved IL1β and GAPDH as loading control upon overnight treatment with nigericin, valinomycin (10µM), salinomycin (10µM) or monensin (10µM). Data are representative of three independent experiments. Immunoblot representative of three replicate experiments.

**Figure 6.**
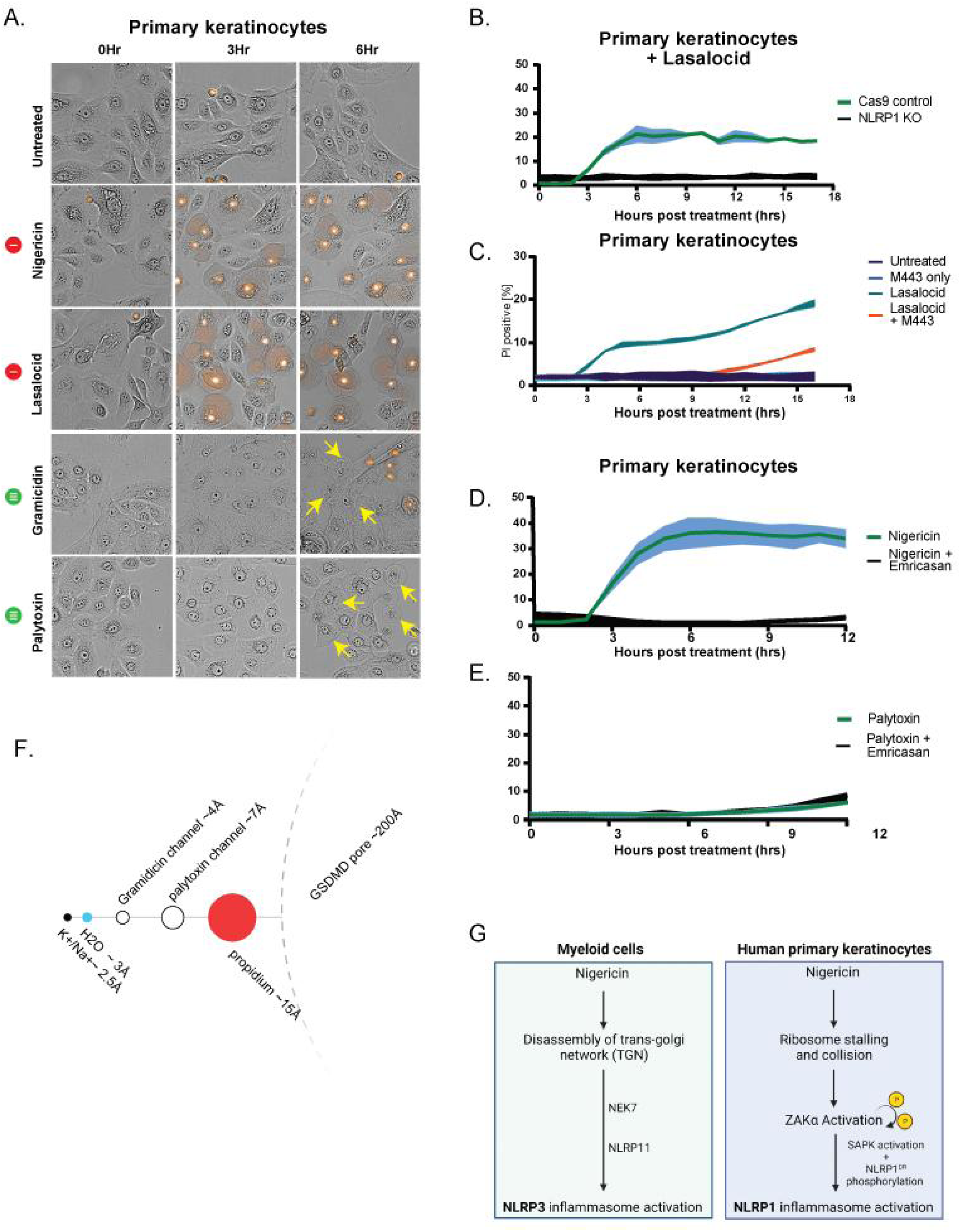
A. Brightfield and PI inclusion images of primary keratinocytes upon nigericin (5µg/mL), lasalocid (5µM), gramicidin (1µM) or palytoxin (100pM) 0, 3 and 6 hours post treatment. Arrows indicate lysed cells that are PI negative. Representative data are shown for n = 3. B. Quantification of the percentage of PI positive Cas9 control and NLRP1 KO primary keratinocytes upon lasalocid (5µM) treatment. Cells were imaged at 15 minute intervals for 18 hours. C. Quantification of the percentage of PI positive primary keratinocytes with or without pre-treatment with M443 (1µM) before lasalocid treatment (5µM).Cells were imaged at 15 minute intervals for 18 hours. D. Quantification of the percentage of PI positive primary keratinocytes with or without pre-treatment with emricasan (5µM) before nigericin (5µg/mL) E. Quantification of the percentage of PI positive primary keratinocytes with or without pre-treatment with emricasan (5µM) before palytoxin (100pM) treatment.Cells were imaged at 15 minute intervals for 18 hours. F. Comparison of the inner diameters of the respective channels/pore to the diameters of H2O and ions. Diagram drawn to scale. G. Summary of how nigericin activates NLRP3 and NLRP1 in distinct human cell types.

**Table 1.**
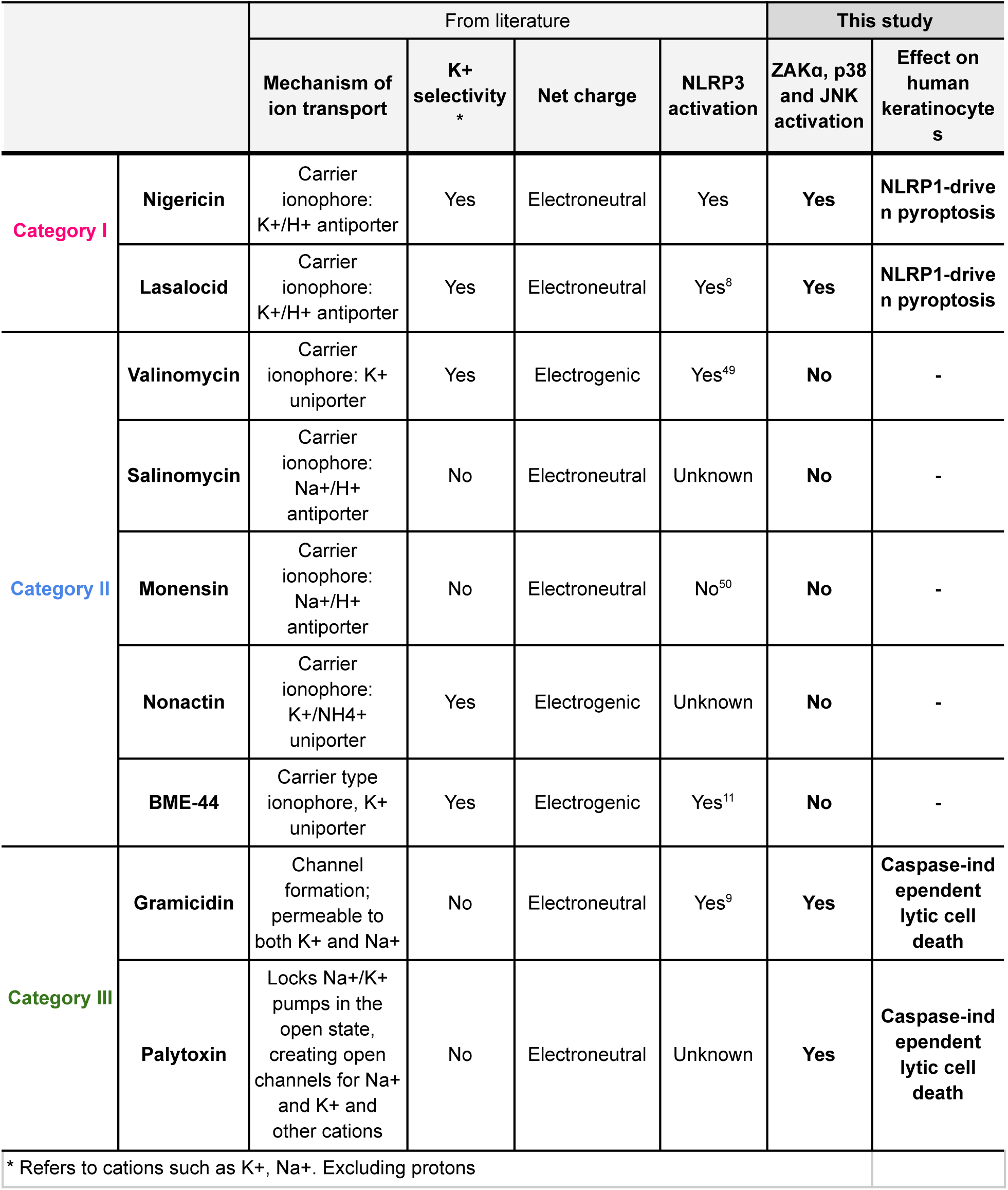
Properties of molecules tested and their effects on RSR and NLRP1 inflammasome

|  |  | From literature |  |  |  | This study |  |
| --- | --- | --- | --- | --- | --- | --- | --- |
| | | Mechanism of ion transport | K<sup>+</sup> selectivity * | Net charge | NLRP3 activation | ZAKα, p38 and JNK activation | Effect on human keratinocytes |
| Category I | Nigericin | Carrier ionophore: K<sup>+</sup>/H<sup>+</sup> antiporter | Yes | Electroneutral | Yes | Yes | NLRP1-driven pyroptosis |
|  | Lasalocid | Carrier ionophore: K<sup>+</sup>/H<sup>+</sup> antiporter | Yes | Electroneutral | Yes<sup>8</sup> | Yes | NLRP1-driven pyroptosis |
| Category II | Valinomycin | Carrier ionophore: K<sup>+</sup> uniporter | Yes | Electrogenic | Yes<sup>49</sup> | No | - |
|  | Salinomycin | Carrier ionophore: Na<sup>+</sup>/H<sup>+</sup> antiporter | No | Electroneutral | Unknown | No | - |
|  | Monensin | Carrier ionophore: Na<sup>+</sup>/H<sup>+</sup> antiporter | No | Electroneutral | No<sup>50</sup> | No | - |
|  | Nonactin | Carrier ionophore: K<sup>+</sup>/NH<sub>4</sub><sup>+</sup> uniporter | Yes | Electrogenic | Unknown | No | - |
|  | BME-44 | Carrier type ionophore, K<sup>+</sup> uniporter | Yes | Electrogenic | Yes<sup>11</sup> | No | - |
| Category III | Gramicidin | Channel formation; permeable to both K<sup>+</sup> and Na<sup>+</sup> | No | Electroneutral | Yes<sup>9</sup> | Yes | Caspase-independent lytic cell death |
|  | Palytoxin | Locks Na<sup>+</sup>/K<sup>+</sup> pumps in the open state, creating open channels for Na<sup>+</sup> and K<sup>+</sup> and other cations | No | Electroneutral | Unknown | Yes | Caspase-independent lytic cell death |
| * Refers to cations such as K<sup>+</sup>, Na<sup>+</sup>. Excluding protons |  |  |  |  |  |  |  |

Category II consists of ionophores that transport K+ and/or other cations in an electrogenic fashion without the ‘antiport’ of a cation or the ‘synport’ of an anion (Table 1). Although they can permeabilize the plasma membrane towards K+, they cannot deplete the intracellular K+ concentration as completely as nigericin due to the creation of extra charge. This likely explains why none of these ionophores have obvious effect on protein synthesis, ribosome elongation (Figure 5B, 5D, S12A), ZAKɑ or SAPK phosphorylation (Figure 5A). Some Category II ionophores, such as valinomycin, are cytotoxic but do not lead to bona fide NLRP1 inflammasome activation, as measured by IL-1β p17 cleavage (Figure 5D, 5E). The fact that Category II molecules have been shown to activate NLRP3, but are unable to activate RSR or NLRP1, suggests that NLRP3 might be more sensitive to K+ efflux than ZAKɑ/NLRP1.

The behaviors of Category III molecules, gramicidin and palytoxin shed further light on how K+ efflux activates RSR. Gramicidin forms oligomeric channels in the plasma membrane that are permeable to many cations ^38^. Palytoxin is a marine toxin that locks the Na+/K+ ATPase pump into the open state, essentially converting it into a cation channel ^39, 40^. Despite their different modes of action, both molecules induce rapid outflow of K+ and inflow Na+, and thus the complete dissipation of the membrane potential. If our hypothesis linking K+ efflux and RSR were true, both molecules would be predicted to be strong activators of ribosome stalling and RSR, similar to Category I molecules. Indeed, gramicidin and palytoxin induced rapid loss of puromycin incorporation and abolished ribosome run-off (Figure 5C). This is accompanied by SAPK phosphorylation and ZAKɑ autophosphorylation (Figure 5A). ZAKɑ KO partially but significantly reduced p38 and JNK phosphorylation induced by gramicidin and palytoxin (Figure S12B), suggesting other stress responsive pathways are also activated, in addition to ZAKɑ-driven RSR. Supplementation of extracellular potassium in palytoxin-treated cells rescued global protein translation and dampened p38 and JNK phosphorylation (Figure 12D). However, in contrast to nigericin and lasalocid, neither gramicidin nor palytoxin activates the NLRP1 inflammasome (Figure 5D, 6A, S14A) or the hyperphosphorylation of the NLRP1 linker domain that is characteristic of ZAKɑ-driven NLRP1 activation (Figure S12C). We hypothesized that other cell death pathways might compete or inhibit NLRP1-driven pyroptosis in palytoxin and gramicidin treated cells. By tracking the fate of cells via live imaging, we found that both palytoxin and gramicidin cause rapid membrane swelling but these ’ballooned’ cells, unlike pyroptotic cells, only take up PI with a significant delay (>30 mins) (Figure 6B). This is in sharp contrast to ANS- and Nigericin-treated cells, where PI inclusion invariably occurs within 5 mins, or sometimes precedes visible membrane ballooning. Crucially, gramicidin- and palytoxin-induced lysis cannot be inhibited by the pan-caspase inhibitor, emricasan (Figure 6D-E, S15A). These observations suggest that palytoxin and gramicidin do not cause pyroptosis, but instead kill the cells via direct osmotic lysis, likely mediated by water influx. This notion is supported by the physical dimensions of these channels. Gramicidin channels are ∼4Å in diameter ^41^, whereas palytoxin locks the Na+/K+ ATPase pump into an open channel with an inner diameter of ∼7Å ^39^. Thus, both channels are large enough to allow free passage of monovalent cations as well as water molecules (∼2Å) (Figure 6F). The channel/pore dimensions also explain why gramicidin- and palytoxin lysed cells do not become immediately permeable to PI as seen in nigericin- or lasalocid-treated cells: propidium ions are ∼15Å in diameter and are therefore too large to pass through the palytoxin- and gramicidin-induced channels ^42^. For comparison, GSDMD pores that perforate pyroptotic cells are >200Å in diameter. We postulate that rapid osmotic lysis interferes with NLRP1 phosphorylation and inflammasome activation, although additional experiments are required to confirm this hypothesis and decipher the biochemical details.

In summary, these results demonstrate that NLRP1 and NLRP3 have differing sensitivities toward membrane disrupting agents. While NLRP3 can activated by multiple triggers that cause K+ efflux, ZAKɑ-driven RSR and the NLRP1 inflammasome activation likely requires a higher threshold of K+ efflux. Thus, only electroneural, K+ selective ionophores such as nigericin and lasalocid acid activate NLRP1 in a ZAKɑ-dependent manner in primary human epithelial cells. Channel-forming molecules such as palytoxin and gramicidin, which cause electroneutral membrane permeabilization via simultaneous K+ efflux and Na+ influx, also activate ZAKɑ-driven RSR, but the concomitant osmolytic lysis likely interferes with NLRP1 activation.

## DISCUSSION

In this study, we demonstrate that cytosolic K+ efflux elicited by ionophores such as nigericin and lasalocid acid is a common trigger for two different inflammasome sensors: NLRP1 in primary human epithelial cells and NLRP3 in hematopoietic cells. In multiple types of primary epithelial cells, nigericin causes rapid K+ efflux-dependent cessation of protein synthesis and ribosome stalling, which in turns activates ZAKɑ-driven RSR and SAPKs. The combined action of ZAKɑ and p38 kinase then drives NLRP1 activation through the hyperphosphorylation of its linker domain.

It is well known that human NLRP1 and NLRP3 have non-overlapping co-factors, agonist repertoire as well as tissue distribution ^12, 16–18, 23, 24, 43^. While NLRP3 is mostly expressed in cells of hematopoietic origin, human NLRP1 is preferentially expressed in epithelial cells in the skin and airway. In fact, none of the cell types or cell lines we have tested so far encode functional NLRP1 and NLRP3 simultaneously. Furthermore, NLRP1 and NLRP3 mutations cause distinct human diseases ^44^. Our results here, however, demonstrate that NLRP1 and NLRP3 could sense a common trigger: the loss of potassium ion homeostasis. That said, only a small subset of K+ ionophores, exemplified by nigericin and lasalocid acid are able to activate both NLRP1 and NLRP3. By analyzing a panel of diverse ionophores, we infer that ZAKɑ-driven NLRP1 activation likely requires a greater degree of K+ efflux than NLRP3, which could only be achieved by electroneutral K+ ionophores (Category I in Table 1). By contrast, electrogenic K+ ionophores, such as valinomycin, which have been shown to activate NLRP3 in multiple studies, are unable to activate RSR or NLRP1 (Category II in Table 1), likely because the extent to K+ flux is thermodynamically constrained by the net separation of charge. Supporting this notion, earlier studies have shown that nigericin and lasalocid acid are also more efficient than valinomycin at activating NLRP3 ^8^. We also showed that more drastic membrane perforation caused by gramicidin channels and palytoxin-hijacked Na+/K+ pumps (Category III in Table 1) also activate ZAKɑ-driven RSR and SAPK activation, but they fail to activate the NLRP1-driven pyroptosis, likely because of interference from direct osmolysis caused by sodium and water influx. This finding shed light on the toxicology of these molecules. For instance, palytoxin exposure has been shown to cause respiratory distress as well as skin lesions ^45^. It is likely that rapid induction of ZAKɑ and osmotic lysis of epithelial cells both contribute to the pathogenesis of palytoxin poisoning.

In summary, we show here that K+ efflux is a broad-acting danger signal that activates RSR and commits multiple human cell types to pyroptotic cell death via distinct inflammasome sensors. Considering that most of the known ionophores, including nigericin, are microbial metabolites, we speculate that the shared ability of NLRP1 and NLRP3 to sense K+ efflux allow them to collaboratively defend multiple organs against certain uncharacterized pathogen/pathogen(s). In this regard, it is interesting to note that regulated membrane depolarization has recently been found to play an active role in CRISPR-mediated antiphage immunity in bacteria ^46^. Our results also strengthen the recently described connection between ZAKɑ-dependent RSR and inflammation and gives credence to the recently proposed ‘HAMP’ hypothesis ^47^, which posits that the human innate immune system monitors key homeostatic processes (e.g. ribosome elongation) in order to respond to a wide variety of seemingly unrelated triggers using a limited number of sensor proteins.

## ACKNOWLEDGEMENTS

The authors are grateful for useful discussion and scientific advice from all members of the Zhong lab. We would like to thank Dr. Esther Koh (NOBIC) for assistance with image acquisition and analysis. We also thank Dr. Tomasz Próchnicki and Prof. Eicke Latz (University of Bonn, Germany) for sharing their experience and knowledge on NLRP3. Illustrations used in this manuscript were created with BioRender.com. Work from F.L.Z. ’s lab is funded by the National Research Foundation Fellowship, Singapore (NRF-NRFF11-2019-0006) and Nanyang Assistant Professorship (NAP). Work in the Bekker-Jensen lab was supported by the European Research Council (ERC) under the European Union’s Horizon 2020 research and innovation program (grant agreement 863911 - PHYRIST).Work from E. Meunier’s group was supported by the European Research Council (ERC) (ERC Starting Grant INFLAME 804249) and the French National Agency for Research (ANR, PSICOPAK).

## METHODS

### Cell culture and chemicals

Immortalized human keratinocytes (N/TERT) were provided by H. Rheinwald (MTA). These cells were cultured in Keratinocyte Serum Free Media (Gibco, 17005042), supplemented with 294.4 ng/l human recombinant Epidermal Growth Factor (EGF) (Gibco, 10450-013), 25 mg/l Bovine Pituitary Extract (Gibco, 13028-014) and 300µM of CaCl2 (Kanto Chemicals, 07058-00). 293Ts (ATCC, #CRL-3216), A549 (Invivogen, a549-ascg-nlrp1), THP-1 (Invivogen, thpd-nfis), U937 (ATCC, #CRL-1593.2*), U20S (ATCC, #HTB-96) . All cell lines used in this study underwent routine mycoplasma testing with MycoGuard (Genecopoeia #LT07-118). Primary human keratinocytes were derived from the foreskin of two donors of Malay ethnicity and obtained with informed consent from the Asian Skin Biobank (ASB) (https://www.a-star.edu.sg/sris/technology-platforms/asian-skin-biobank). pHNECs (Sigma Aldrich, SCCE016).

The following drugs and chemicals were used as part of this study: Nigericin (Sigma, N7143), Anisomycin (ANS, MCE, #HY18982), Talabostat (VbP, MCE, #HY-13233), MCC950 (MCE, HY-12815A), ATP (Thermofisher Scientific, R0441), Belnacasan (MCE, HY-13205), Emricasan (MCE, HY-10396), Potassium chloride (Sigma Aldrich, P9541), BAPTA (MCE, HY-100168), Harringtonine (HTN, MCE, HY-N0862), Puromycin (PURO, Sigma, #P9620), M443 (MCE, HY-112274), Neflamapimod (p38i) (MCE, HY-10328), Bentamapimod (JNKi) (MCE, HY-14761), Valinomycin (MCE, HY-N6693), Salinomycin (MCE, HY-15597), Monensin (MCE, HY-N0150), Gramicidin (MCE, HY-P0163), Palytoxin (Fujifilm Wako, 165-26141), Nonactin (Sigma Aldrich, N2286), BME-44 (Sigma Aldrich 60397), Lasalocid (MCE, HY-B1071), Emetine (EME, MCE, HY-B1479B). Compound 6p is a kind gift from X. Lu (Jinan University, China)

### Cytokine analysis

Human IL-1β enzyme linked immunosorbent assay (ELISA) kit (BD, #557953) and human IL-18 ELISA kit (R&D, DY318-05) were used in accordance with the manufacturer’s protocols to measure secreted cytokines.

### Immunoblotting and antibodies

Cells that were attached to wells were lysed directly with 1x Laemmli buffer (100µl of 1x LB for 200,000 cells in a 12 well plate). To visualize cleaved GSDMD, cells that were floating in the supernatant (floaters) were separated after collection of media supernatant and lysed directly with 1x LB (30µl of 1x LB from a 12 well plate). All lysates were boiled at 95°C for 5 minutes before loading. For the detection of cleaved GSDMD, lysates from the floater fraction and the attached cell fraction were combined at a 1:1 ratio (15µl floater + 15µl attached cell lysates) prior loading. For visualizing IL-1β in the media, harvested supernatant was concentrated using filtered centrifugation (Merck, Amicon Ultra, #UFC500396). All protein samples were run using immunoblotting and the Chemidoc Imaging System (Bio-rad) was used for visualization. PhosTag SDS-PAGE were carried out using homemade 8% or 10% SDS-PAGE gel, supplemented with Phos-tag Acrylamide (Wako Chemicals, AAL-107) to a final concentration of 30μM and manganese chloride(II) (Sigma-Aldrich, #63535) to 60μM. Cells were also directly lysed in 1x LB and boiled before loading into the Phos-tag gel. Upon run completion, the polyacrylamide gel was washed in transfer buffer supplemented with 10mM EDTA twice and a final wash was done without EDTA. Each wash was done for 10 minutes. All SDS-PAGE gels were blotted onto 0.45μm PVDF membranes (Bio-rad) and blocked with 3% milk for 1 hour prior incubation with the respective primary antibodies overnight at 4℃. The following day, the corresponding secondary antibodies were used and blots were incubated for 1 hour at room temperature. The following antibodies were used in this study: IL1β p17 specific (Cell Signaling Technology, #83186S), pro–IL-1β (R&D Systems, AF-401-NA), GFP (Abcam, ab290), GAPDH (Santa Cruz Biotechnology, #sc-47724), GSDMDC1 (Novus Biologicals, NBP2-33422), GSDME (Abcam, ab215191), Total p38 (Abcam, ab32142), phospho-p38 (Cell Signaling Technology, #4511), Total JNK (Cell Signaling Technology #9252), phospho-JNK (Cell Signaling Technology, #4668), ZAK (Bethyl Laboratories, A301-993A), puromycin (Merck Millipore, MABE343), MK2 (Abcam, ab32567), EIF2A-P (Cell Signaling Technology, #3398), NLRP1 (Biolegend, #9F9B12), ASC (Adipogen, AL177). All horseradish peroxidase (HRP)-conjugated secondary antibodies were purchased from Jackson Immunoresearch (goat anti-mouse IgG: 115-035-166; goat anti-rabbit IgG: 111-035-144).

### Propidium iodide inclusion assay

Primary keratinocytes of various genotypes were seeded in a black 12-well plate (Cellvis, P12-1.5P) at a density of 180,000 cells per well. The following day, cells were treated with the respective chemicals and stained with 0.5 µg/mL of propidium iodide (PI, Abcam #ab14083) before the plate was loaded into a high content screening microscope (Perkin Elmer Operetta CLS imaging system, NTU Optical Bio-Imaging Centre in Nanyang Technological University, Singapore) for overnight imaging. Brightfield and fluorescence (PI channel, 536 nm/ 617 nm) were captured every 15 mins. Images were then stored and analyzed using the Harmony software (Version 6). Using data acquired from 5 fields of view per well with three wells per treatment, the ratio of PI positive cells over the total number of cells was calculated. The number of live cells per field was calculated using digital phase contrast images which allows for the identification of cell borders. The number of PI positive stained cells were identified through the PI channel and counted as PI positive cells.

### Harringtonine ribosome run off assay

Human primary keratinocytes were seeded at a cell density of 80,000 cells/well in a 24 well plate. The next day, cells were pre-treated with 10µM of emricasan for 30 minutes followed by treatment with the relevant ionophores or anisomycin for 3 hours. The respective samples were treated with 2µg/mL of harringtonine at staggered timepoints of 5 min, 1 min and 0 min (equivalent to mock untreated). Cells were then pulsed with a final concentration of 10 µg/mL of puromycin at the same time in all wells for 10 mins. Following puromycin treatment, the supernatant was discarded and cells were lysed directly in 1x Laemmli buffer. Immunoblotting of samples using an anti-puromycin antibody was done to measure the amount of nascent peptides with puromycin incorporated, which reflects the rate of elongation after treatment.

### Plasmid and preparation of lentiviral stock

N/TERT NLRP1 KO + NLRP1^DR^-GFP, N/TERT ZAKα KO cells were previously described ^27^. Constitutive lentiviral expression was performed using pCDH vector constructs (System Biosciences) and packaged using third-generation packaging plasmids. pSpCas9(BB)-2A-Puro (PX459) V2.0, which was used for generating ZAK KO A549 cells, was a gift from Feng Zhang (Addgene plasmid # 62988 ; http://n2t.net/addgene:62988 ; RRID:Addgene_62988). pPBbsr-JNK KTR-mCherry (Addgene plasmid # 115493 ; http://n2t.net/addgene:115493 ; RRID:Addgene_115493) and pNJP (Addgene plasmid # 115494 ; http://n2t.net/addgene:115494 ; RRID:Addgene_115494) were gifts from Kazuhiro Aoki. pLentiPGK-Blast-DEST-JNKKTRmRuby2 was a gift from Markus Covert (Addgene plasmid # 59154 ; http://n2t.net/addgene:59154 ; RRID:Addgene_59154). These plasmids were used to generate p38 and JNK Kinase Translocation Reporter(KTR) cells.

### CRISPR-Cas9 Knockout

NLRP1 KO and MAP3K20 (ZAKα) KO N/TERT keratinocytes were made and described in detail previously ^27^.MAP3K20 (ZAKα) KO human primary keratinocytes were generated using lentiviral Cas9 and guide RNA plasmid (LentiCRISPR-V2, Addgene plasmid #52961) using the following guides: sg1 (TGTATGGTTATGGAACCGAG), sg4 (TGCATGGACGGAAGACGATG). NLRP1 KO human primary keratinocytes were generated using lentiviral transduction with the following guide: sg1 (GATAGCCCGAGTGCATCGG). A549-ASC-GFP-NLRP1 MAP3K20 (ZAKα) KO cells were generated using transfection of pSpCas9 plasmids with sgRNA into cells, followed by a diphtheria toxin selection method. In brief, A549 cells were seeded with a density of 100,000 cells in a 12WP. Cells were transfected using FuGENE HD on the next day with two separate pSpCas9-2A-puro plasmids, one encoding MAP3K20 guide 4(TGCATGGACGGAAGACGATG) and another encoding HBEGF guide 10 (CACCTCTCTCCATGGTAACC), in a 4:1 ratio. 2 days post transfection, cells were treated with 20ng/ml of diphtheria toxin(DT) for selection. Only cells that have the DT entry receptor HBEGF knocked out would survive DT treatment and would likely have undergone gene-editing in the MAP3K20 gene. Knockout efficiency was tested by immunoblot. Alternatively, Sanger sequencing of genomic DNA and overall editing efficiency were determined using the Synthego ICE tool (Synthego Performance Analysis, ICE Analysis. 2019. v2.0. Synthego, https://ice.synthego.com/<u>#/</u>).

### In-vitro Translation Assay

In-vitro translation assay was performed using T_N_T® Quick Coupled Transcription/Translation System (Promega #L1170) following the manufacturer’s recommendations. Briefly, luciferase mRNA (TriLink) was used to measure translational regulation by nigericin. In total, 1µg/mL of mRNA were added to 20ul of TnT mastermix supplemented with methionine. The reaction was incubated for 60 mins at 30℃. 1µl of reaction was added to 25µl of luciferase assay reagent (Promega) and luminescence activity was measured in each well.

### Generating p38 and JNK KTR

The KTR system was adapted from published studies ^48, 49^. For clarity, p38 ‘KTR’ refers to GFP-MK2, while the JNK ‘KTR’ refers to the original design consisting of a JNK substrate motif. Both KTR constructs were cloned into pCDH lentiviral expression vectors and expressed via stable transduction in N/TERT cells.

### Microscopy

All microscopy images were acquired at NTU Optical Bio-Imaging Centre (NOBIC) imaging facilities at LKCMedicine. Live cell imaging was carried out with Operetta CLS High Content Analysis System (PerkinElmer) and confocal microscopy was carried out with Olympus FV3000 (Olympus Life Sciences).

### Polysome profiling

Cells were exposed to nigericin (5 mg/mL) at the indicated time. Following treatment, cells were lysed in a lysis buffer containing 20 mM Hepes pH 7.5, 100 mM NaCl, 5 mM MgCl_2_, 100 mg/ml digitonin, 100 mg/ml cycloheximide, 1X protease inhibitor cocktail (Sigma, #P8340) and 200 U NxGen RNase inhibitor (Lucigen, #30281). Extracts were incubated on ice for 5 min prior to centrifugation at 17,000 g for 5 min at 4 °C. After adding calcium chloride to a final concentration of 1 mM, lysates were optionally digested with 500 U micrococcal nuclease (MNase) (New England Biolabs, #M0247) for 30 min at 22 °C. Digestion was quenched by adding 2 mM EGTA. Equivalent amounts of lysate (250 mg of undigested RNA or 350-400 mg of MNase-digested RNA) were run through 15-50% sucrose gradients. Gradients were centrifuged at 38,000 rpm in a Sorvall TH64.1 rotor for two hours at 4 °C. The gradients were passed through an ISCO density gradient fractionation system with continuous monitoring of the absorbance at 254 nm.

### Sucrose cushion

Cells were exposed to nigericin (5mg/mL) at the indicated time. Following treatment, cells were lysed (15 mM Tris, pH 7.5, 0.5% NP40, 6 mM MgCl2, 300 mM NaCl, RiboLock) and centrifuged at 12000 g, 4°C, 10 min. The supernatant was carefully layered onto a sucrose cushion (30% sucrose in 20 mM Tris, pH 7.5, 2 mM MgCl2, 150 mM KCl) and ultra-centrifuged at 38800 rpm for 16 h using Sorvall™ wX+ Ultrafuge and FIBERlite F50L-8x39 rotor. Pellets were resuspended in 100 mM KCl, 5mM MgCl2, 20 mM HEPES, pH 7.6, 1 mM DTT and 10 mM NH4Cl. Purified ribosome fractions were analyzed by western blot

## FIGURES AND LEGENDS

All significance values were calculated based on one-way ANOVA from three biological replicates, with each treatment considered a single replicate. Significance values were indicated as: n.s (non significant), ***P<0.01, ***P<0.001, ****P<0.0001.

---

**Supplementary Figure S1.**
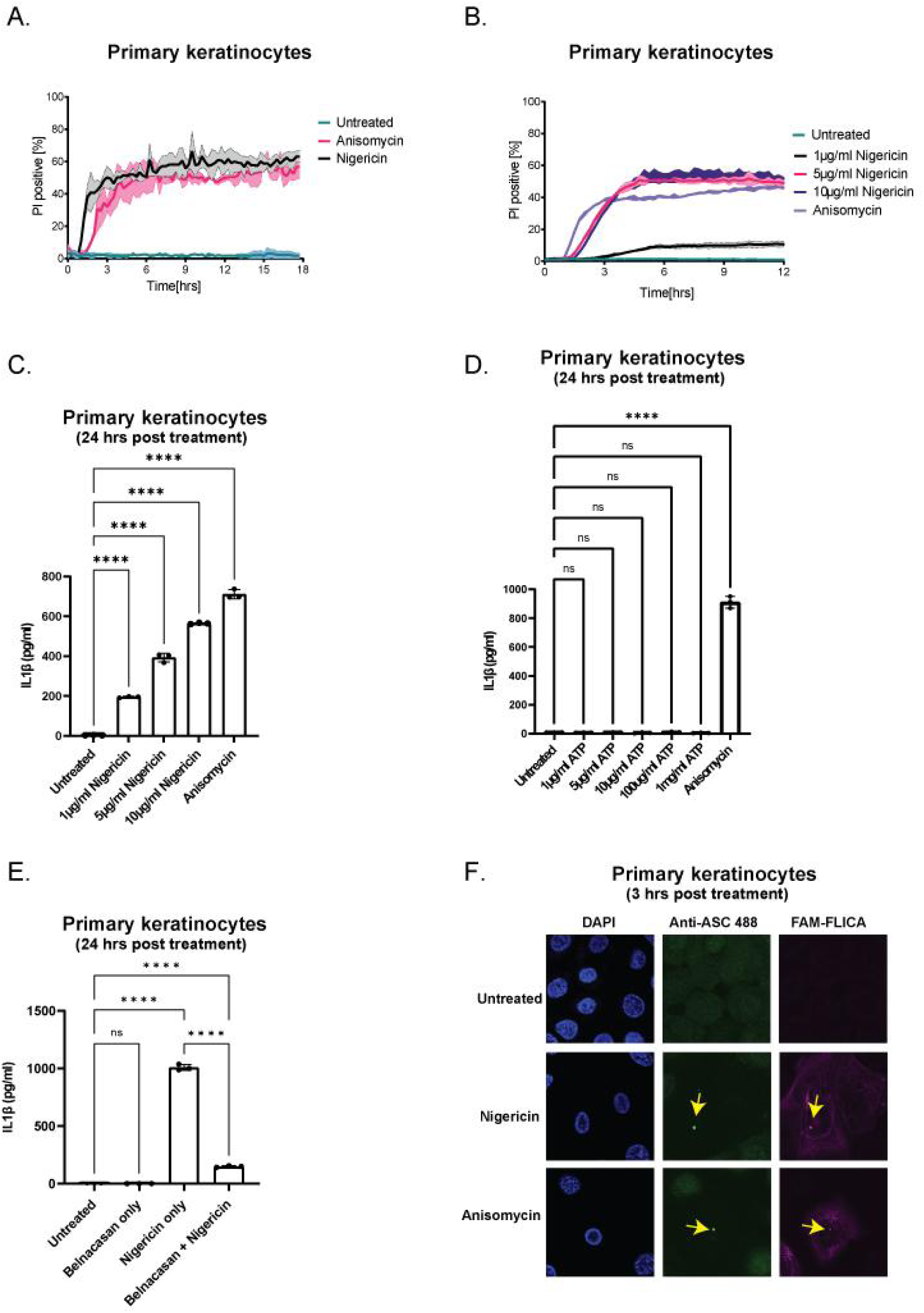
A. Quantification of the percentage of PI positive WT primary keratinocytes after nigericin (5µg/mL) or anisomycin (1µM) treatment. Cells were imaged at 15 minute intervals for 18 hours. B. Quantification of the percentage of PI positive WT primary keratinocytes after nigericin (1µg/mL, 5µg/mL or 10µg/mL) or anisomycin (1µM) treatment. Cells were imaged at 15 minute intervals for 18 hours. C. ELISA showing IL-1β secretion in primary keratinocytes after nigericin (1µg/mL, 5µg/mL or 10µg/mL) or anisomycin (1µM), or D. ATP (1µg/mL, 5µg/mL, 10µg/mL or 100µg/mL), or E. nigericin (5µg/mL) +/- Belnacasan (5µM). Supernatant was harvested 24 hours post treatment. F. Representative confocal images showing ASC (green), Caspase-1 (FAM-FLICA) and DAPI (blue) staining in primary keratinocytes 3 hours post nigericin (5µg/mL) or anisomycin (1µM) treatment.

**Supplementary Figure S2.**
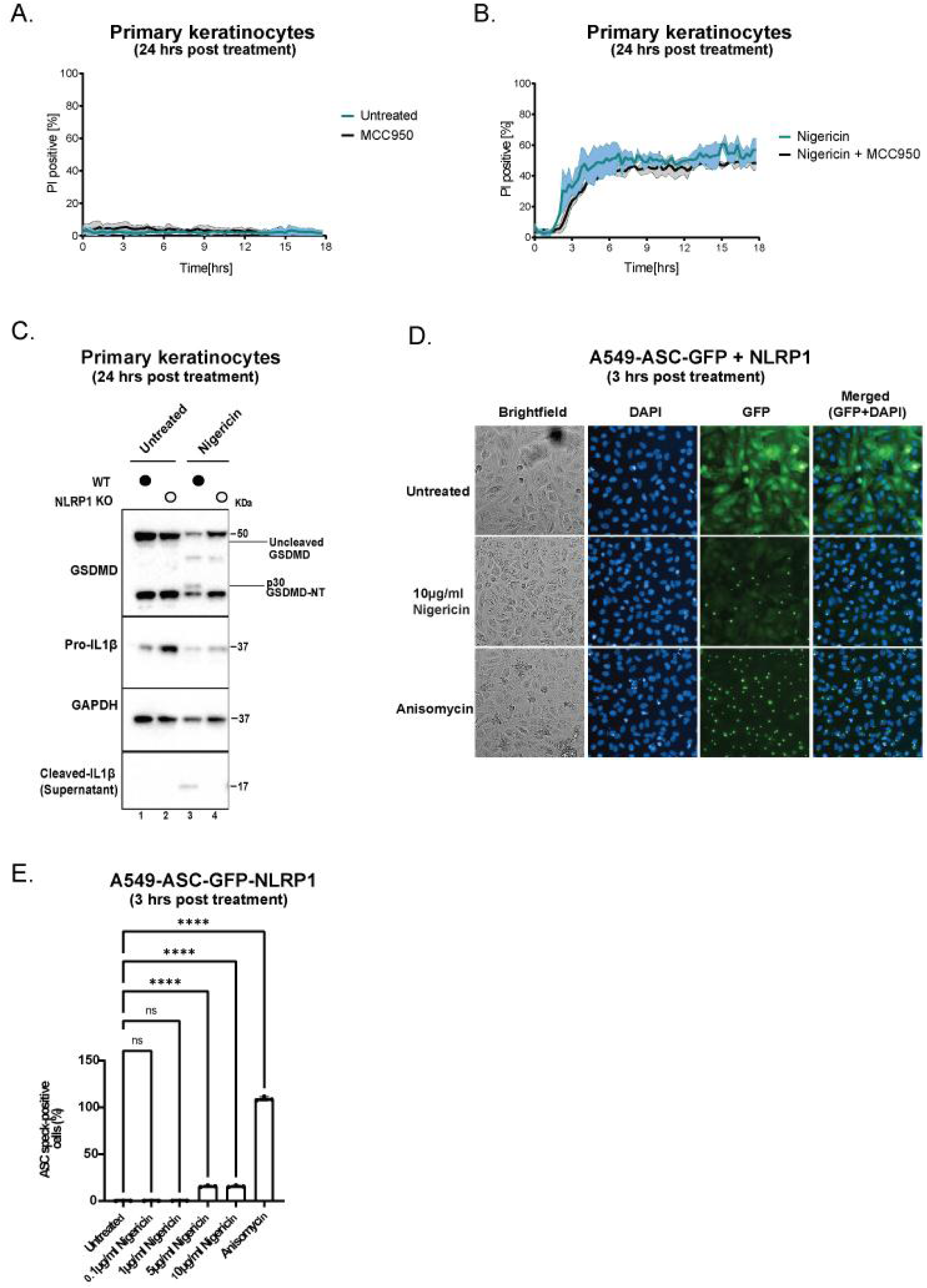
A. Quantification of the percentage of PI positive WT primary keratinocytes with MCC950 (5µM) only or, B. with nigericin (5µg/mL) and nigerinc+MCC950 (5µM). Cells were imaged every 15 minutes for 18 hours. C. Western blot analysis of WT and NLRP1 KO primary keratinocytes showing GSDMD (full length and cleaved), pro-IL1β, cleaved IL1β and GAPDH upon overnight treatment with nigericin (5µg/mL). Immunoblot representative of three replicate experiments. D. Representative images of A549 cells overexpressing ASC-GFP and NLRP1. Images of GFP and DAPI fluorescence were acquired 3 hours post treatment with nigericin (10µg/mL) or anisomycin (1µM) at 20x magnification. E. Percentage of ASC-speck forming cells in A549 overexpressing ASC-GFP and NLRP1 upon 3 hour treatment with nigericin (0.1µg/mL, 1µg/mL, 5µg/mL or 10µg/mL) or anisomycin (1µM).

**Supplementary Figure S3.**
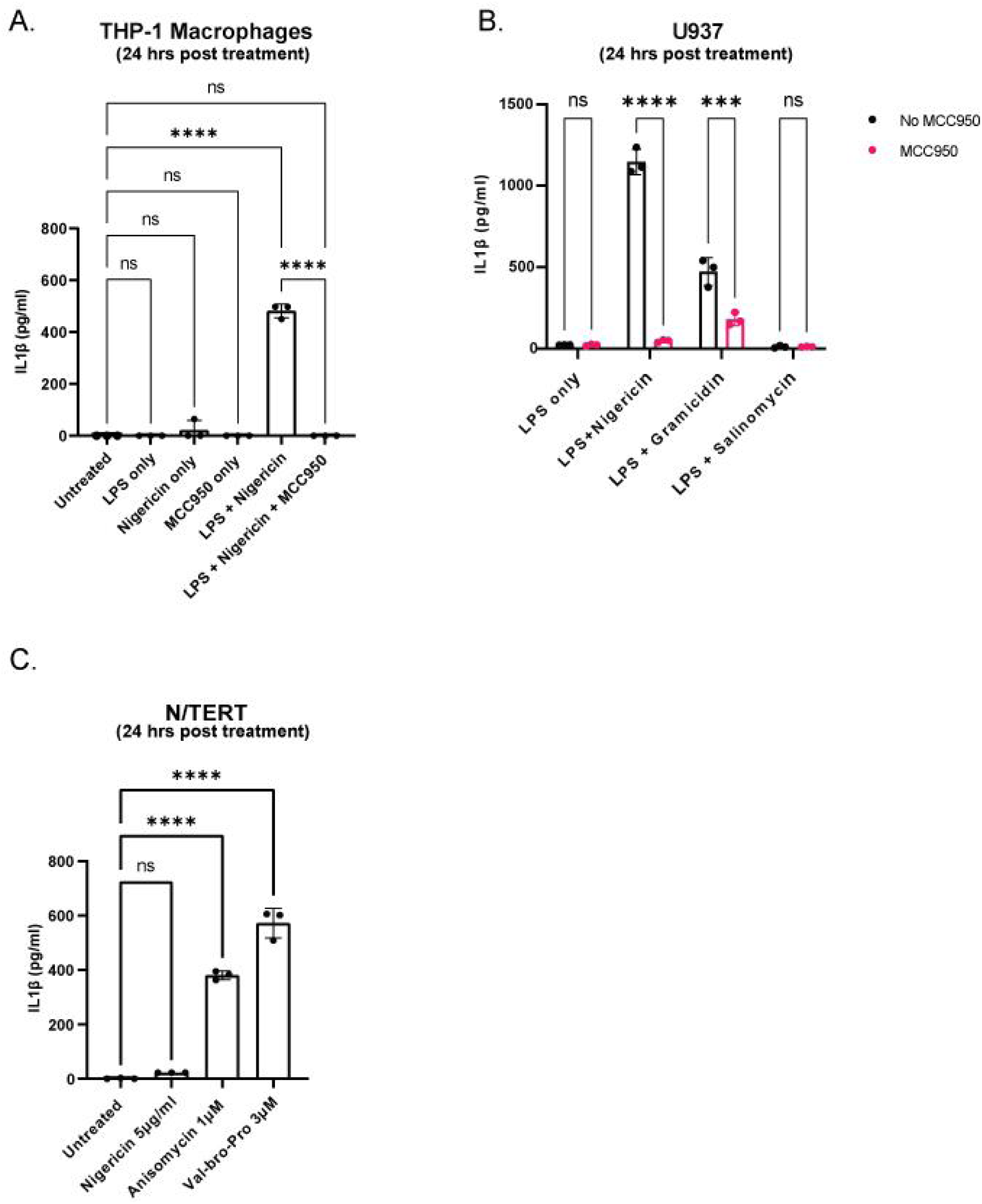
A. ELISA showing IL-1β secretion in PMA-differentiated (200ng/ml overnight) THP-1 macrophages. Differentiated THP-1 macrophages were further stimulated with LPS (1µg/mL) for 3 hours followed by nigericin (5µg/mL) treatment overnight. B. ELISA showing IL-1β secretion in PMA-differentiated (200ng/ml overnight) U937 macrophages. U937 macrophages were stimulated with LPS (100ng/ml) for 3 hours and nigericin (5µg/mL), gramicidin (1µM) or Salinomycin (5µM) overnight with or without MCC950 (5µM). C. ELISA showing IL-1β secretion in N/TERTs after overnight treatment with nigericin (5µg/mL), ANS (1µM) or VbP (3µM).

**Supplementary Figure S4.**
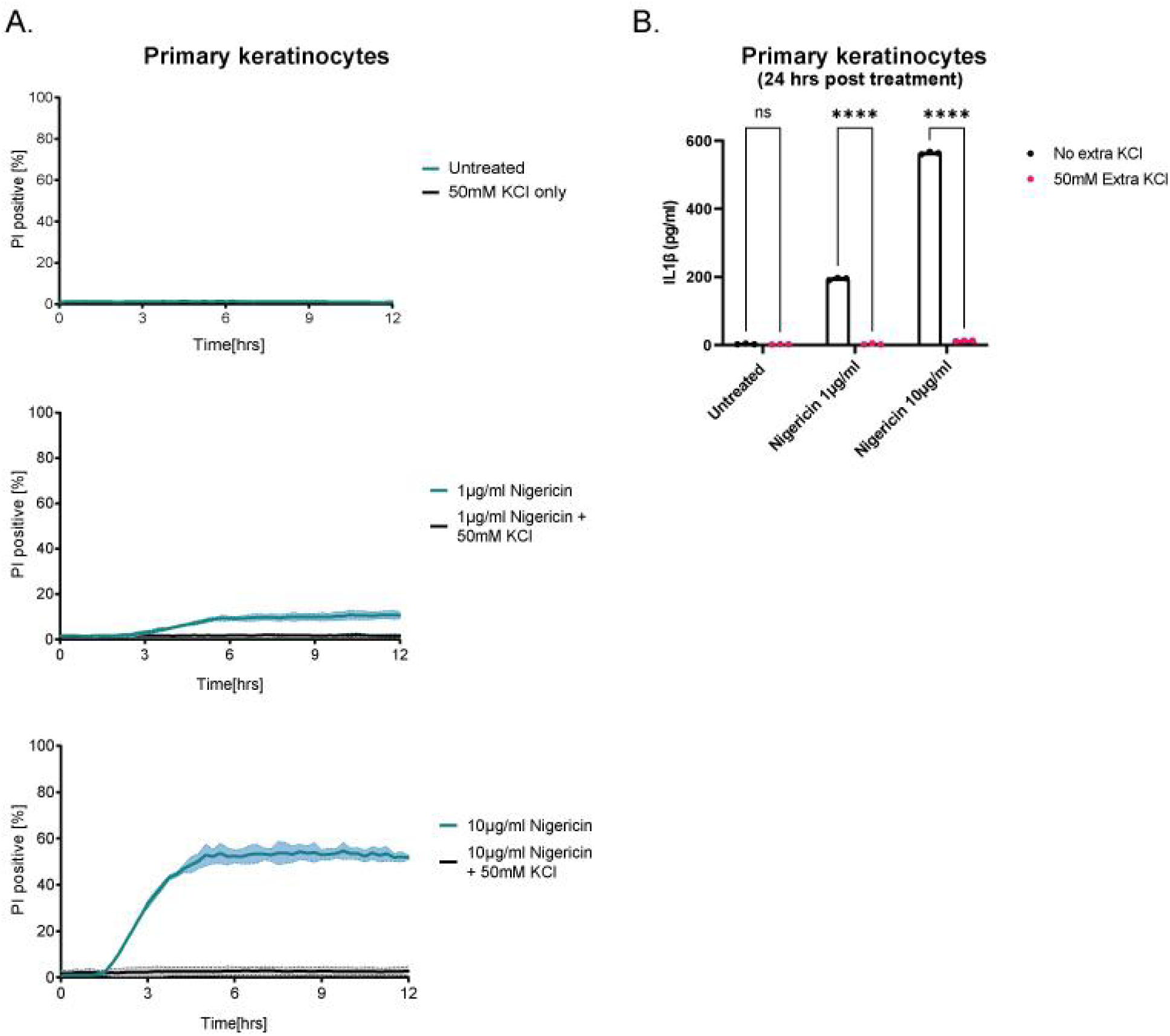
A. Quantification of the percentage of PI positive WT primary keratinocytes treated with KCl only (50mM), nigericin only or nigericin + KCl. Cells were imaged at 15 minute intervals for 12 hours. B. ELISA showing IL-1β secretion from supernatant obtained from (A).

**Supplementary Figure S5.**
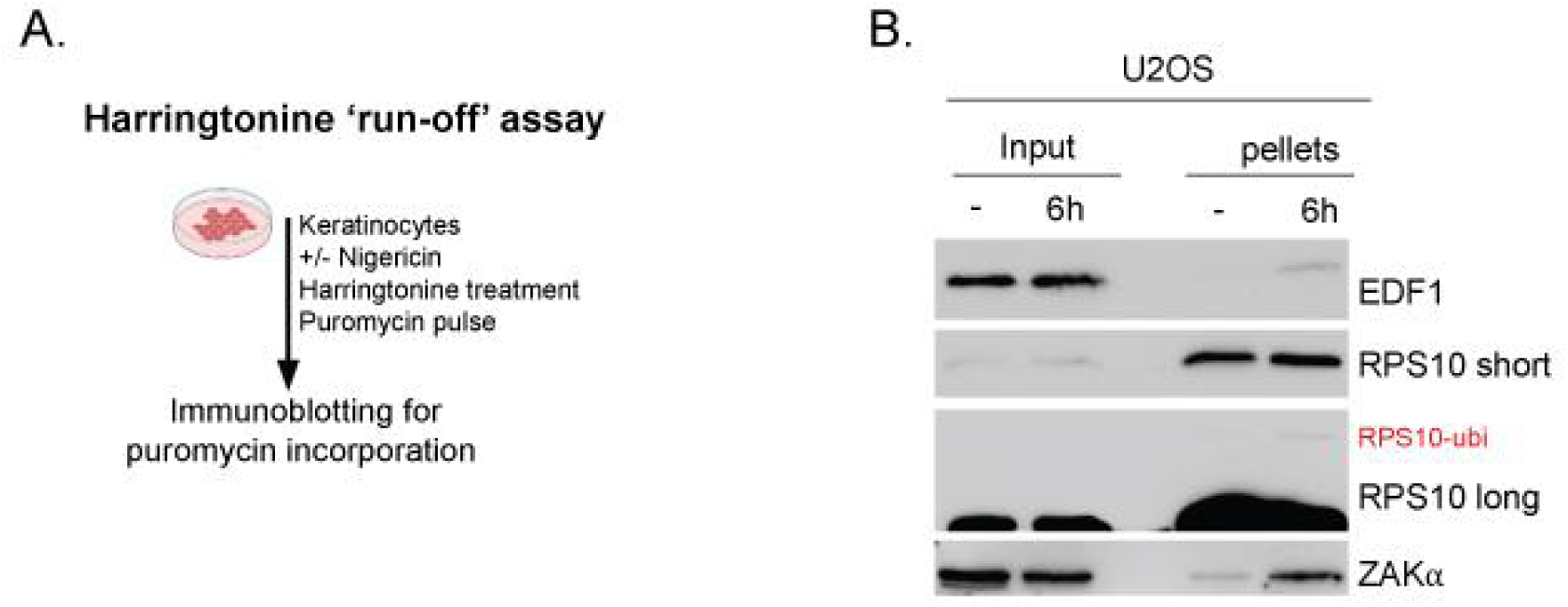
A. The workflow of the harringtonine ‘run-off’ assay B. U2OS cells were treated with nigericin (5mg/mL) for 6 hours and lysates were ultra-centrifuged through a sucrose cushion to collect the ribosome fraction (Pellet). Input lysate and pelleted material were analyzed by western blot.

**Supplementary Figure S6.**
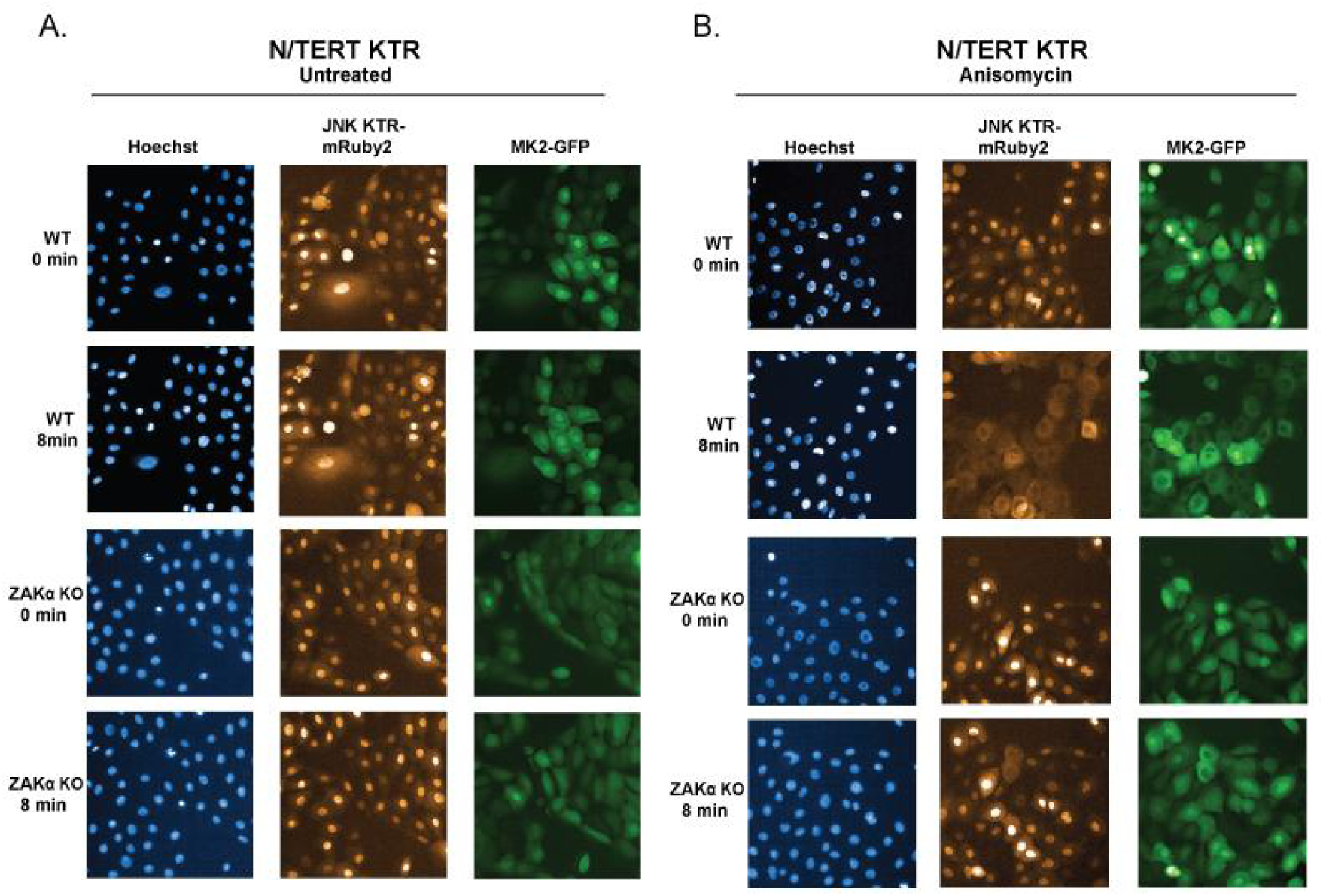
A-B. Representative images of WT N/TERT cells stably expressing MK2-mEGFP and JNK KTR-mRuby2 upon stimulation with anisomycin (1µM). Images were acquired every 4 minutes for 2 hours.

**Supplementary Figure S7.**
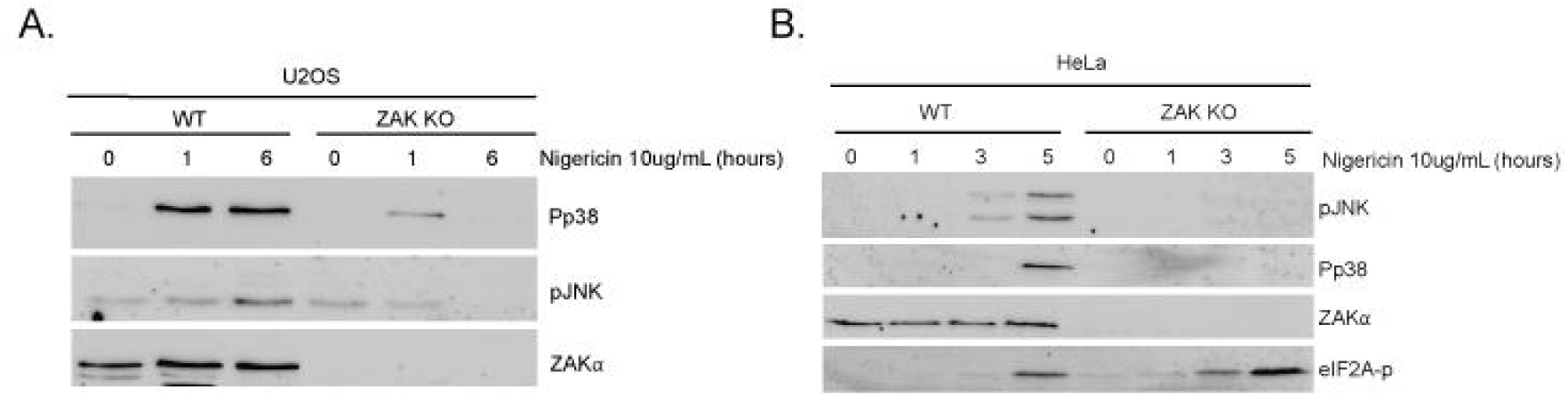
A. Western blot showing the induction of p-p38 and pJNK in WT and ZAK KO U2OS cells. B. Western blot showing the induction of p-p38 and pJNK in WT and ZAK KO HeLa cells.

**Supplementary Figure S8.**
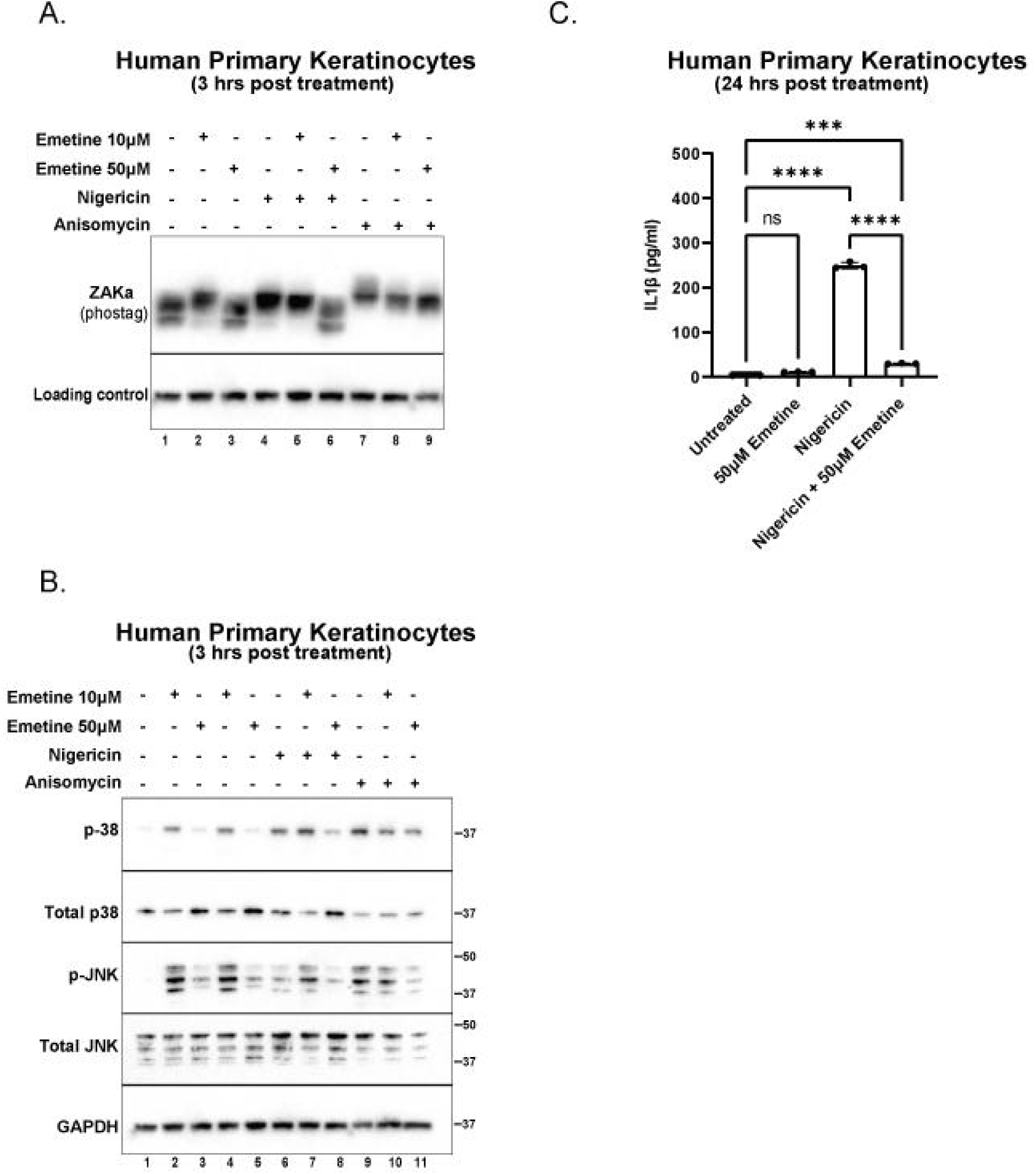
A. Western blot analysis of PhosTag SDS-page (ZAKɑ) in primary keratinocytes pre-treated with emetine (10µM or 50µM for 15 mins) followed by 3 hour treatment with nigericin (5µg/mL) or anisomycin (1µM). Immunoblot representative of three replicate experiments. B. Western blot analysis of p-p38, total p38, p-JNK, total JNK and GAPDH (loading control) in primary keratinocytes following pre-treatment with emetine (10µM or 50µM for 15 mins) and 3 hour treatment with nigericin (5µg/mL) or anisomycin (1µM). Immunoblot representative of three replicate experiments. C. ELISA showing IL-1β secretion from supernatant obtained from primary keratinocytes following pre-treatment with emetine (50µM for 15 mins) and overnight treatment with nigericin (5µg/mL).

**Supplementary Figure S9.**
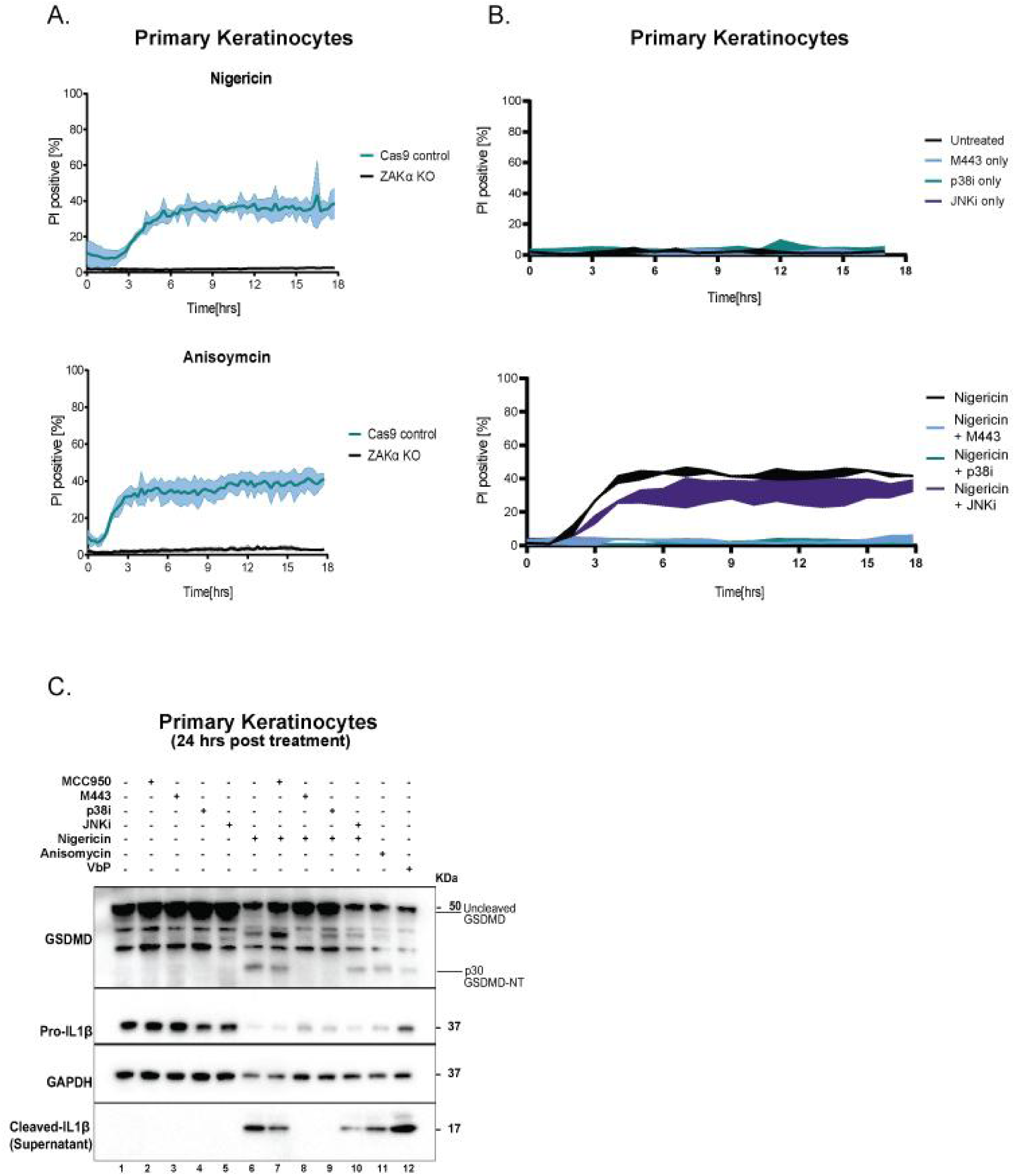
A. Quantification of the percentage of PI positive WT and ZAKɑ KO primary keratinocytes treated with nigericin (5µg/mL) or anisomycin (1µM). Cells were imaged at 15 minutes interval for 18 hours. B. Quantification of the percentage of PI positive WT primary keratinocytes pre-treated (15 min) with various inhibitors p38i (1µM), JNKi (1µM) or M443 (1µM) followed by overnight treatment with nigericin (5µg/mL). Cells were imaged at 15 minute intervals for 18 hours. C. Western blot analysis of primary keratinocytes showing GSDMD (full length and cleaved), pro-IL1β, cleaved IL1β and GAPDH as loading control upon pre-treatment with p38i (1µM), JNKi (1µM) or M443 (1µM) followed by overnight treatment with nigericin (5µg/mL), anisomycin (1µM) or VbP (3µM). Data are representative of three independent experiments. Immunoblot representative of three replicate experiments.

**Supplementary Figure S10.**
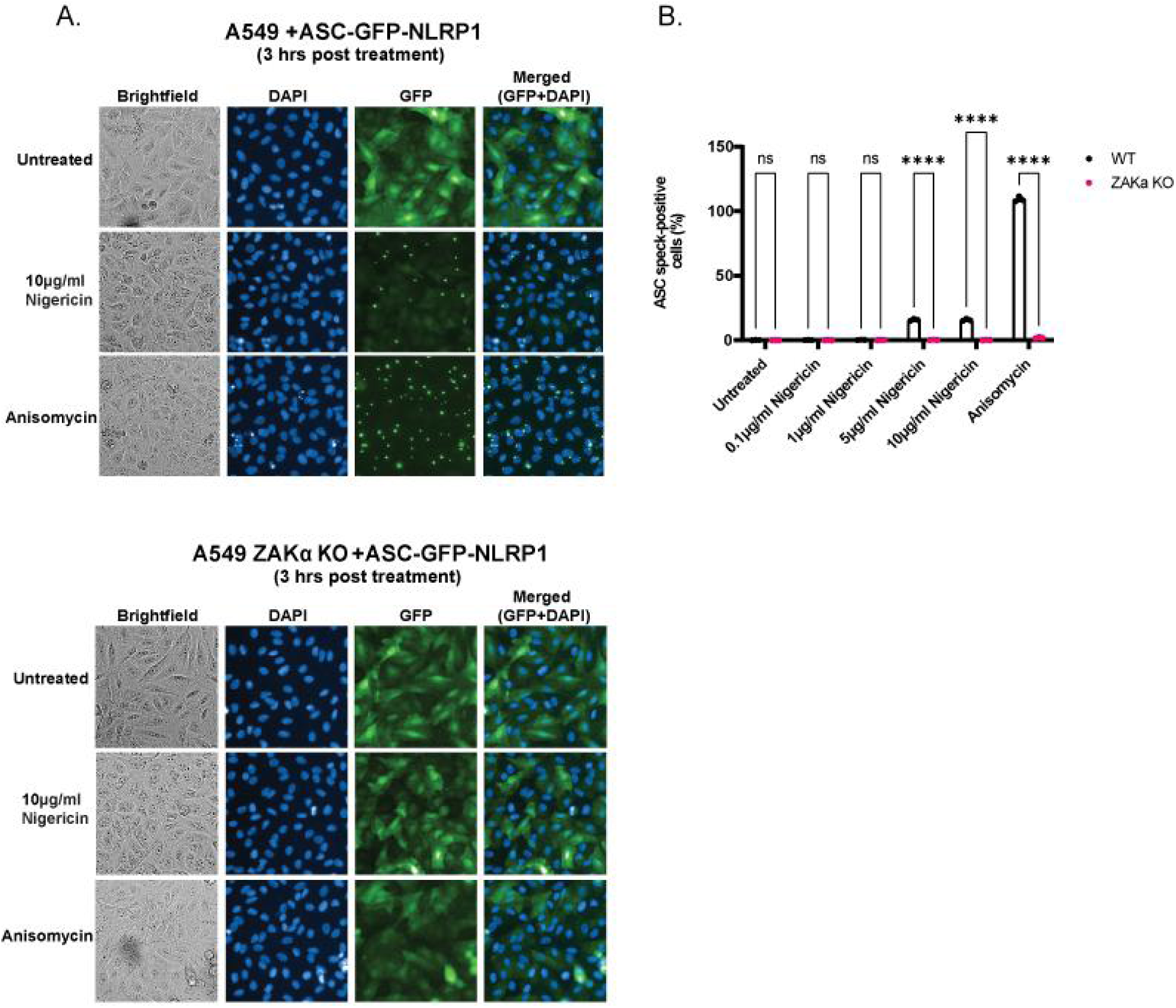
A. Representative images of WT or ZAKɑ KO A549 cells overexpressing ASC-GFP and NLRP1. Images of GFP and DAPI fluorescence were acquired 3 hours post treatment with nigericin (5µg/mL) or anisomycin (1µM) at 20x magnification. B. Quantification of the percentage of ASC-speck forming cells in WT or ZAKɑ KO A549 overexpressing ASC-GFP and NLRP1 in (A).

**Supplementary Figure S11.**
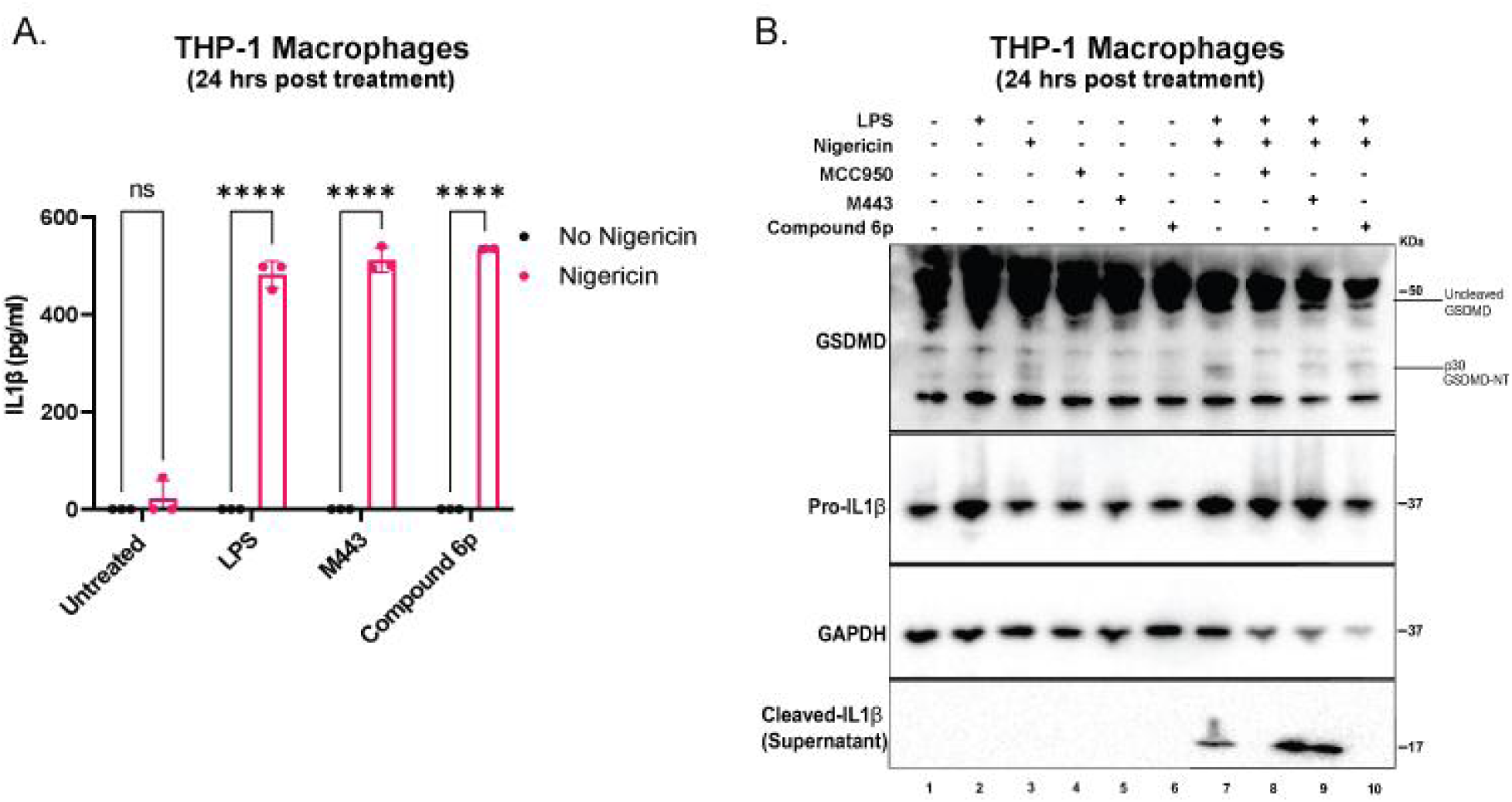
A. ELISA showing IL-1β secretion in PMA-differentiated (200ng/ml overnight) THP-1 macrophages. Differentiated cells were then stimulated with LPS (100ng/ml) for 3 hours followed by pre-treatment with either M443 or compound 6p for 15 mins prior nigericin treatment overnight. B. Western blot analysis of THP-1 macrophages obtained from (A) Immunoblot representative of two replicate experiments.

**Supplementary Figure S12.**
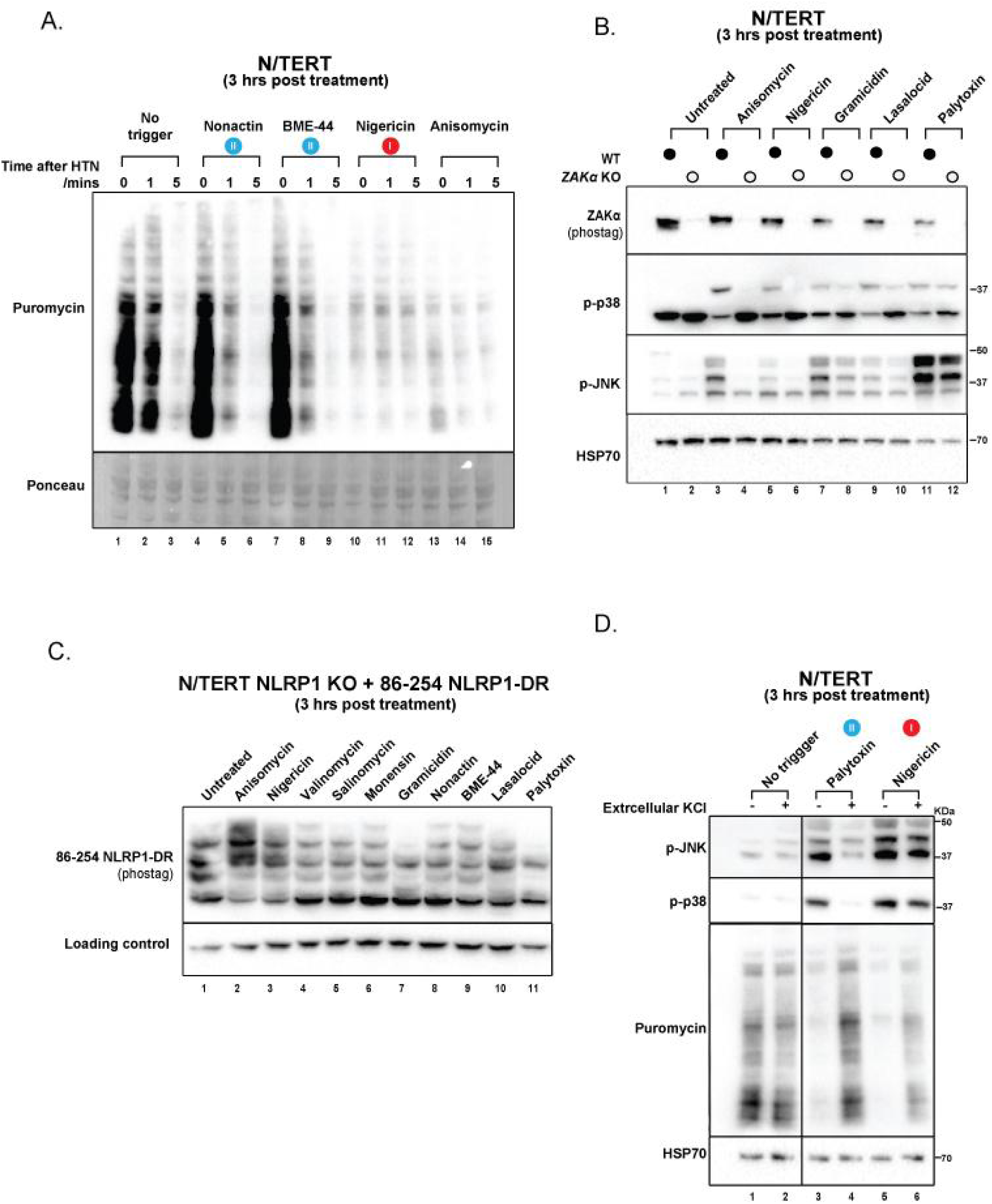
A. Anti-puromycin immunoblots of N/TERT cells subjected to Harringtonine run-off assay. Cells were treated for 3 hours with nonactin (5µM), BME-44 (5µM), nigericin (5µg/mL) or anisomycin (1µM) before harringtonine addition (2µg/mL) for the respective durations. Nascent peptides were then labeled with puromycin (10µg/mL) for 10 mins. Ponceau staining was used as the loading control. Immunoblot representative of three replicate experiments. B. Western blot analysis of PhosTag SDS-page (ZAKɑ) or SDS-page (p-p38, p-JNK and HSP70) of WT or ZAKɑ KO N/TERT cells upon 3 hour treatment with anisomycin (1µM), nigericin (5µg/mL), gramicidin (1µM), lasalocid (5µm) or palytoxin (100pM). Immunoblot representative of three replicate experiments. C. PhosTag SDS-page of NLRP1 KO + NLRP1^DR^ (a.a.86-254)-GFP N/TERT following 3 hours treatment with ANS (1µM), nigericin (5µg/mL), valinomycin (5µM), salinomycin (5µM), monensin (5µM), gramicidin (1µM), nonactin (5µM), BME-44 (5µM), lasalocid (5µM) or palytoxin (100pM). Immunoblot representative of three replicate experiments. D. Western blot analysis of p-p38, p-JNK, puromycin and HSP70 (loading control) in N/TERTs following pre-treatment with 5µM emricasan and 3 hour treatment with palytoxin 100pM or nigericin (5µg/mL) with or without supplementation of 50mM KCl.

**Supplementary Figure S13.**
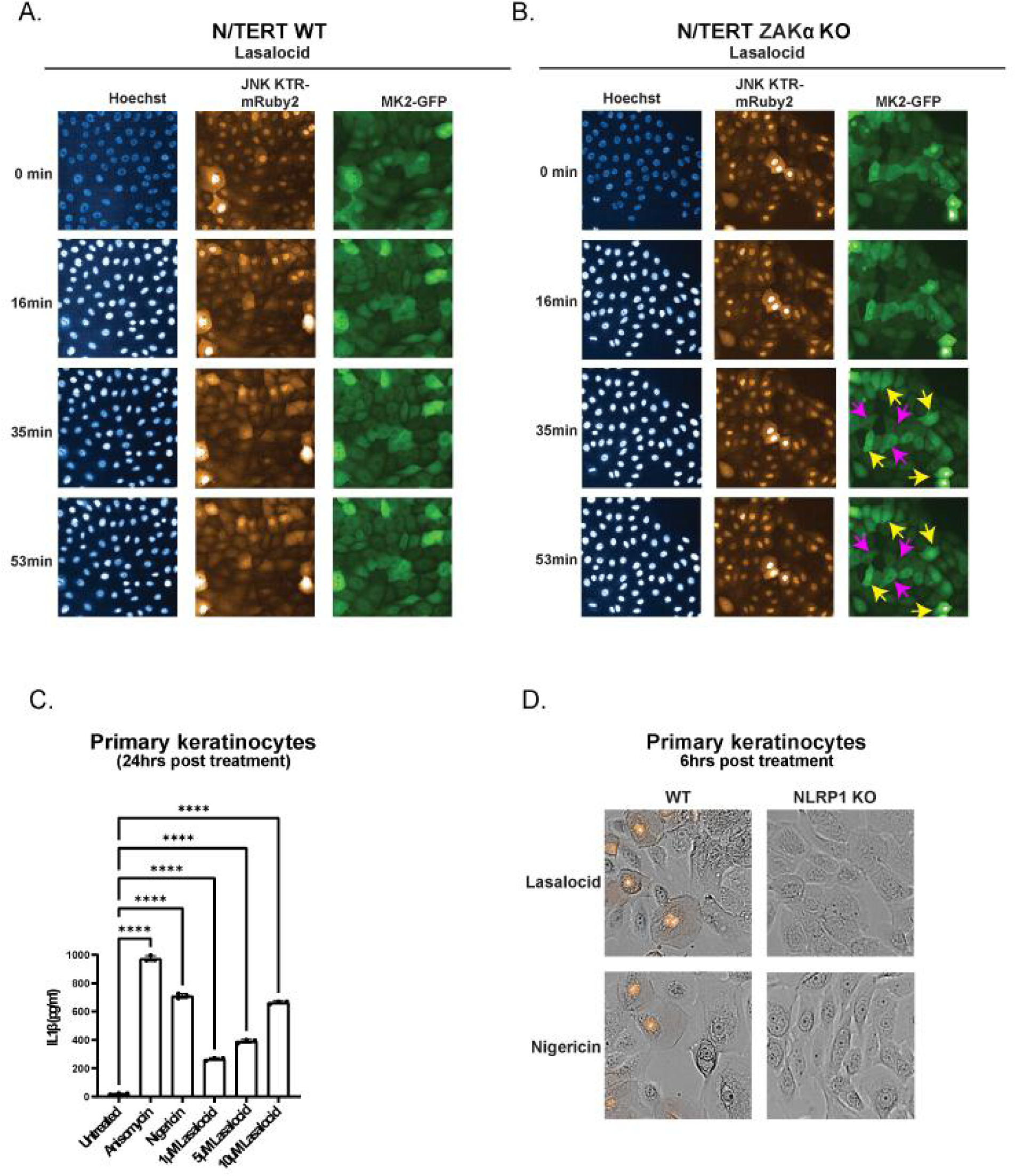
A. Representative images of WT, and B. ZAKɑ KO N/TERT cells stably expressing MK2-mEGFP and JNK KTR-mRuby2 upon stimulation with Lasalocid (5µM). Images were acquired every 8 minutes for 2 hours. Data are representative of three independent experiments. C. ELISA showing IL-1β secretion in primary keratinocytes following overnight treatment with increasing concentrations of lasalocid (1µM, 5µM or 10µM), nigericin (5µg/mL) or anisomycin (1µM). D. Brightfield and PI inclusion images of WT or NLRP1 KO primary keratinocytes upon nigericin (5µg/mL) or lasalocid (5µM) treatment. Images were obtained 6 hours post treatment. Representative data are shown for n = 3.

**Supplementary Figure S14.**
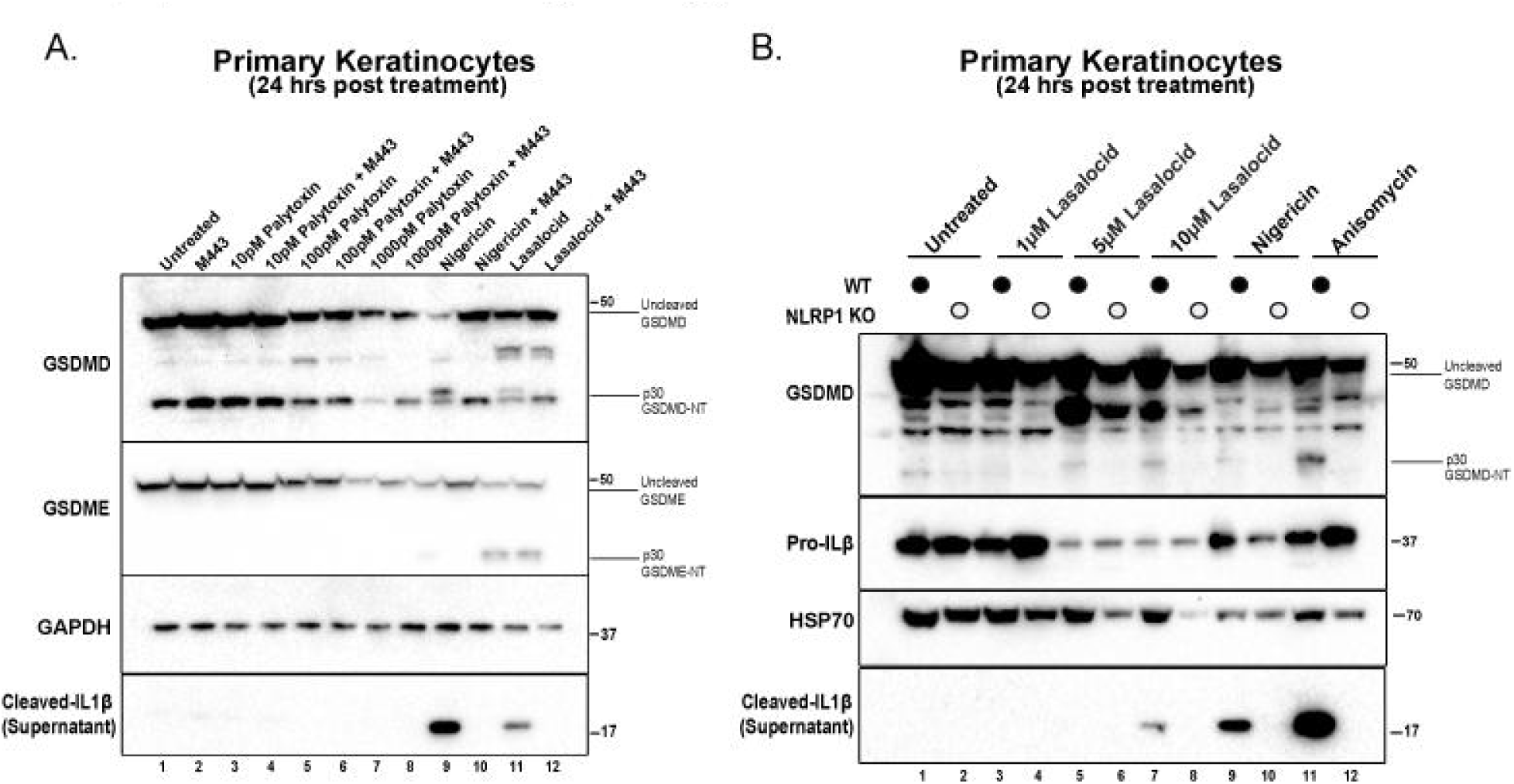
A. Western blot analysis of primary keratinocytes showing GSDMD (full length and cleaved), GSDME (full length and cleaved), pro-IL1β, cleaved IL1β and GAPDH (loading control) upon overnight treatment with palytoxin (10pM, 100pM or 1000pM), nigericin (5µg/mL) or lasalocid (5µM). Pre-treatment with M443 (1µM) for 15 mins was performed for the relevant conditions. Data are representative of three independent experiments. Immunoblot representative of three replicate experiments. B. Western blot analysis of WT or NLRP1 KO primary keratinocytes showing GSDMD (full length and cleaved), pro-IL1β, cleaved IL1β and HSP70 (loading control) upon overnight treatment with increasing concentration of lasalocid (1µM, 5µM or 10µM), nigericin (5µg/mL) or anisomycin (1uM). Data are representative of three independent experiments. Immunoblot representative of three replicate experiments.

**Supplementary Figure S15.**
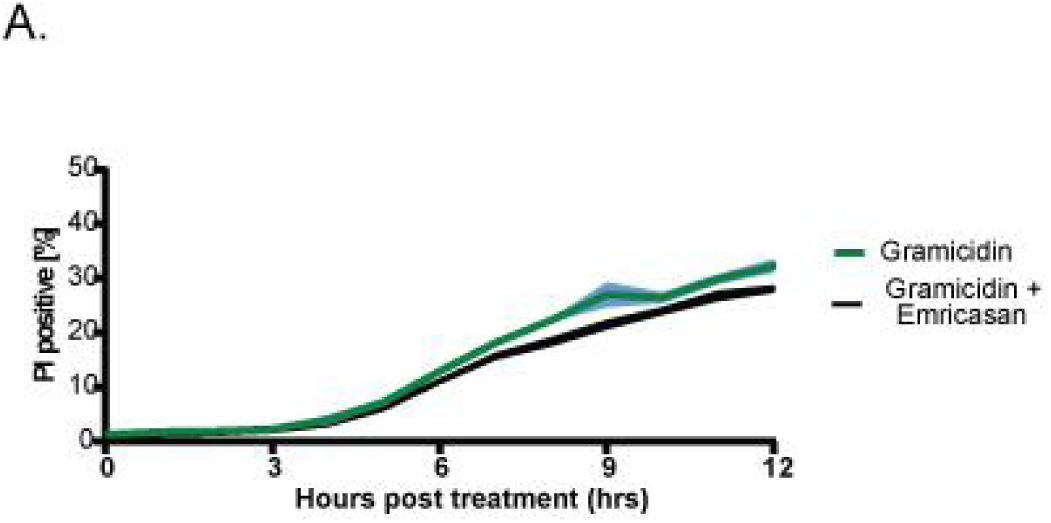
A. Quantification of the percentage of PI positive primary keratinocytes with or without pre-treatment with emricasan (5µM) before gramicidin (1µM) treatment.Cells were imaged at 15 minute intervals for 12 hours.

